# Solvent-buffer effects in molecular dynamics simulations of nucleic acids

**DOI:** 10.64898/2026.07.05.736650

**Authors:** Nainsy Baghel, Pranchal Shrivastava, Rukmankesh Mehra

**Affiliations:** Department of Bioscience and Biomedical Engineering, Indian Institute of Technology Bhilai, Durg – 491002, Chhattisgarh, India; Department of Chemistry, Indian Institute of Technology Bhilai, Durg – 491002, Chhattisgarh, India

**Keywords:** Nucleic acid dynamics, MD simulations, cell size, solvent-buffer distance, solvation shell, ds- versus ss-DNA dynamics

## Abstract

Molecular dynamics simulations of nucleic acids are performed using a solvent-buffer distance of 10 Å between the solute surface and the simulation box boundary. Although this cell size has been extensively explored in protein simulations, its implications for nucleic acid dynamics are not well understood. Nucleic acids are elongated, highly charged, and flexible structures with hydration and dynamical properties distinct from those of proteins and therefore, they may require different solvent-layer considerations in simulations. In this study, we investigated the effect of simulation cell size on nucleic acid dynamics by simulating a 30-base-pair double-helical nucleic acid structure and its two single-stranded forms using solvent-buffer distances of 3, 5, 10, 15, and 20 Å. Smaller cells may impose restricted hydration, molecular crowding, and periodic image interactions. However, larger cells provide solvent space for conformational relaxation. A total of 45 µs of molecular dynamics simulations were performed (3 structures × 5 cell sizes × 3 replicates × 1 µs). Our results show that while the commonly used 10 Å buffer may be sufficient to maintain the stability of the double-stranded nucleic acid, larger cells are required to capture the conformational dynamics of single-stranded structures. In both, increasing the cell size to 15 or 20 Å enables broader conformational sampling. The first hydration shell exhibits reduced crowding in the 20 Å cell, consistent with more relaxed conformations. At larger cell sizes, single-stranded nucleic acids adopt compact, self-associated conformations for stability. Together, this study presents physical insight into how simulation cell size and solvent environment influence nucleic acid dynamics.

## 1 Introduction

Molecular dynamics simulations of macromolecules are commonly performed using a solvent-buffer distance of 10 Å between the solute surface and the simulation box boundary [1–6]. The effect of simulation cell size has been extensively explored in protein simulations, where a 10 Å solvent buffer appears appropriate [2–6]. Protein structures are stabilized by a combination of hydrophobic packing, hydrogen bonding, electrostatic interactions, and interactions with the surrounding hydration shell (**Figure 1a**) [7–10]. Proteins exhibit internal motions, ranging from local side-chain movements to large conformational rearrangements. Their dynamics are often governed by folded-domain stability and collective motions related to function [11, 12]. However, nucleic acids differ substantially from proteins (**Figure 1b**). They are elongated, highly charged, and flexible structures with hydration and dynamical properties distinct from those of proteins [13]. Nucleic acids undergo large-scale motions such as bending, twisting, groove breathing, base flipping, and conformational transitions between different structural forms [14, 15]. Their stability is influenced by hydration networks around the phosphate backbone, sugar groups, and grooves [16, 17]. Therefore, the conformational stability of nucleic acids may require different number of surrounding water layers compared with proteins.

**FIGURE 1.**
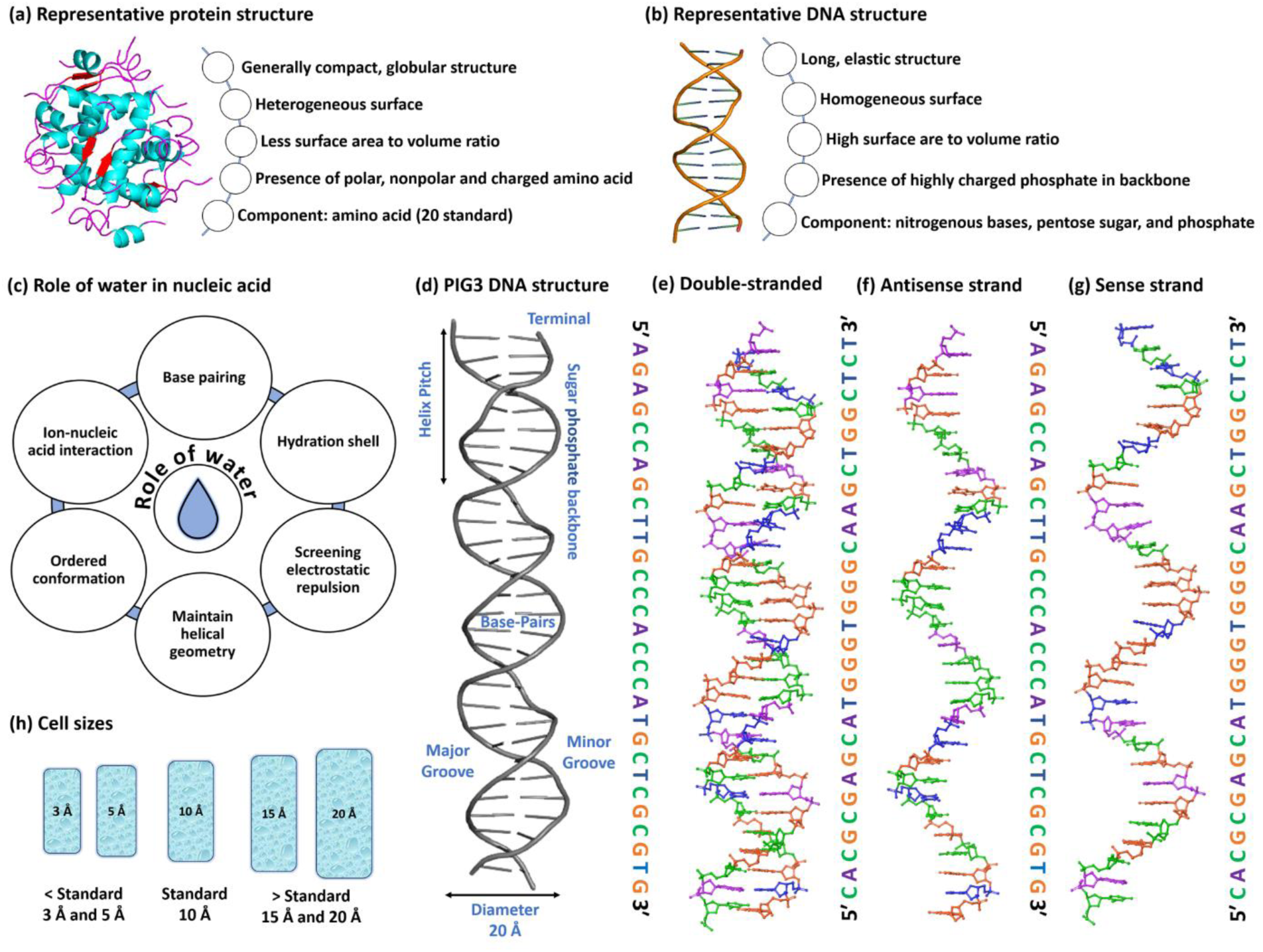
Schematic overview of the molecular systems investigated in this study. **(a)** Representative protein structure used as a structural context. **(b)** Representative double helix structure. **(c)** Role of water molecules in stabilizing nucleic acids. **(d)** Crystal structure of the PIG3 DNA retrieved from the PDB structure 7EDS. **(e)** Double-stranded DNA (dimer) configuration. **(f)** Antisense strand (monomer 1) configuration. **(g)** Sense strand (monomer 2) configuration. **(h)** Schematic illustration of the five periodic simulation cell sizes (3, 5, 10, 15, and 20 Å) used for MD simulations.

Hydration plays a central role in nucleic acid folding and stability (**Figure 1c**). In nucleic acids, water is not just a passive solvent but participates directly in maintaining structure, screening electrostatic repulsion, and enabling conformational flexibility [17–19]. The negatively charged phosphate backbone produces strong electrostatic repulsions, which can destabilize folded structure. Water molecules, together with counterions such as Na^+^, K^+^, and Mg^2+^, reduce this repulsion and allow the nucleic acid chain to adopt ordered conformations [20, 21]. Ordered water molecules around the phosphate backbone, sugar groups, and grooves help maintain helical geometry and can influence sequence-dependent stability [22, 23]. In addition, water layers regulate ion-nucleic acid interactions. Many ions interact with nucleic acids through intervening water molecules rather than by direct binding to the phosphate backbone or bases [24]. These water-mediated ion interactions are essential for stabilizing folded nucleic acid structures and reducing backbone repulsion. Inadequate hydration in simulations may therefore artificially restrict conformational dynamics, alter ion distributions, or destabilize folded structures. Therefore, adequate solvent buffering is necessary for accurately describing nucleic acid stability and dynamics in molecular simulations.

In this study, we systematically investigated how simulation cell size affects the structural and dynamical properties of nucleic acids. Because double-stranded and single-stranded DNA are expected to behave differently during simulations, both forms were considered (**Figure 1d–g**). Single-stranded DNA is particularly suitable for probing cell-size effects because it is expected to sample a broader conformational landscape on the microsecond timescale, which may be difficult to encounter for the stable double-helical structure [25, 26]. Therefore, we performed molecular dynamics (MD) simulations of a 30-base-pair double-stranded DNA system, and its two corresponding single-stranded forms separately using solvent-buffer distances of 3, 5, 10, 15, and 20 Å (**Figure 1h**).The smaller 3 and 5 Å solvent buffers are expected to enhance confinement and possible periodic image interactions, while the larger 15 and 20 Å buffers provide greater solvent space for conformational relaxation [2]. However, increasing the cell size also increases the computational cost significantly because of the larger number of water molecules, ions, and non-bonded interactions.

The aim of this study is to examine: (i) the effect of restricted versus extended solvent buffering; (ii) the effect of possible periodic image interactions; (iii) differences between double-stranded and single-stranded DNA dynamics; (iv) the role of surface and bulk water layers in modulating DNA structure; and (v) the solvent-buffer distance required to adequately sample nucleic acid dynamics.

## 2 Methods

### 2.1 System preparation

The nucleic acid structure of the PIG3 sense and antisense strands was selected from the Protein Data Bank (PDB) structure 7EDS (**Figure 1d**) [27]. This structure contains 30 nucleotides per strand in complex with human p53 protein and was determined at a resolution of 1.77 Å. The PIG3 duplex adopts an approximately B-DNA conformation, which is the predominant DNA form under physiological conditions. The 30-base-pair duplex contains approximately three complete helical turns and provides a sufficiently long template for analyzing nucleic acid structural dynamics.

For system preparation, the p53 protein and other heteroatoms were removed to obtain the isolated double-helical PIG3 DNA structure (**Figure 1d**). From this duplex structure, the sense and antisense strands were also extracted and considered separately (**Figure 1e–g**). The systems were prepared at pH 7.4 using Schrodinger [28]. Hydrogen bonding networks were optimized, and local energy minimization was performed to remove steric clashes.

### 2.2 Molecular dynamics simulations

To evaluate the effect of simulation cell size, the DNA systems were prepared using five solvent-buffer distances: 3, 5, 10, 15, and 20 Å (**Table S1**). The solvent-buffer distance was defined as the minimum distance between the nucleic acid surface and the simulation box boundary in all directions. The smaller buffers of 3 Å and 5 Å were included to assess the possible effects of restricted hydration and periodic image interactions. The 10 Å buffer represented the commonly used setup in biomolecular simulations, whereas the larger 15 and 20 Å buffers were used to provide extended solvent environments and to examine the role of water molecules at increasing distances from the nucleic acid.

In addition to the double-helical structure, the two individual strands were simulated separately as single-stranded systems, namely the antisense (monomer 1) and sense (monomer 2) strands, each containing 30 nucleotides. This ensured the ordered evaluation of the effects on single strands in comparison to the double strands. Because the two strands are complementary, they provide a paired dataset for comparing their dynamics in isolated single-stranded and duplex forms.

In total, 15 systems were prepared, comprising five cell sizes for each of the three DNA systems: double-stranded DNA, antisense strand, and sense strands. Each system was solvated in an orthorhombic box using TIP3P water. Counterions were added to neutralize the system, followed by additional Na^+^ and Cl^-^ ions to maintain 0.15 M salt concentration. Periodic boundary conditions were applied. (**Table S2**) summarizes the system setup, including the number of water molecules, atoms, and ions.

Each system was energy minimized for 100 picoseconds using a combination of steepest descent and Broyden-Fletcher-Goldfarb-Shanno methods [29]. Molecular dynamics simulations were performed in the NPT ensemble at 300 K and 1.01325 bar using a 2 fs time-step. Temperature and pressure were maintained using a Nose-Hoover chain thermostat [30] and a Martyna-Tobias-Klein barostat [31], respectively. Long-range electrostatic interactions were treated using the particle mesh Ewald method with a cutoff of 9.0 Å [32, 33].

Each system was simulated for 1 µs in triplicate, for a total simulation time of 45 µs (1 µs × 3 runs × 15 systems) across all systems. All simulations were performed in Desmond using OPLS4 force field [34]. OPLS4 includes refined parameters for nucleic acids and has been validated against experimental and quantum-mechanical reference data, making it suitable for the present study [35, 36].

## 3 Results and discussion

### 3.1 Analysis of nucleotide content in each system

The PIG3 DNA structure and sequence present in PDB ID 7EDS were analyzed prior to the MD simulations. The duplex contains 30 nucleotides per strand, corresponding to approximately three helical turns. The distance spanning ten consecutive base pairs, corresponding to one helical pitch, ranged from 33.40 to 34.11 Å (**Table S3**). The maximum inter-strand width ranged from 19.68 to 20.36 Å (**Table S4**). These structural parameters are consistent with an initial B-DNA conformation of nucleic acid [37].

The double-helical structure followed Chargaff base-pairing rule[38]. Each single strand, that is the antisense strand and sense strand, contained 16.67% adenine and 16.67% thymine. However, the guanine and cytosine contents differed between the two complementary strands. The antisense strand contained 40.00% cytosine and 26.67% guanine, while the sense strand contained 26.67% cytosine and 40.00% guanine (**Figure 1e–g)**. Despite this strand-specific difference, the overall GC content was 66.67% in each single strand and in the double-helical system (**Table S5**).

### 3.2 Larger solvent buffers allow broader conformational sampling

To evaluate the effect of solvent-buffer distance on nucleic acid dynamics, the structure-function relations of all 15 nucleic acid systems were analyzed side by side. The conformational stability of each system was primarily evaluated from the complete 1 µs trajectories using backbone root-mean-square deviation (RMSD), radius of gyration (R_g_) and solvent accessible surface area (SASA).

For the double-helical structure, the backbone RMSD showed greater variation in larger cells (**Figures 2a**, **S1a** and **Table S6**). In the smaller 3 and 5 Å cells, the RMSD variation was comparatively lower, with standard deviations of 0.61 and 0.78 Å, respectively. This reduced variation may arise from the restricted solvent space available for nucleic acid motion. The average RMSD of the three replicate simulations for the 3, 5, 10, 15 and 20 Å cells were 5.02 ± 0.61, 4.99 ± 0.78, 5.08 ± 0.94, 5.08 ± 1.01, and 5.11 ± 1.01 Å, respectively. Although the average RMSD values were similar across all cell sizes, the fluctuations increased in the larger cells. This suggests that larger solvent buffers allow greater conformational freedom, whereas 3 and 5 Å cells may restrict molecular motion.

**FIGURE 2.**
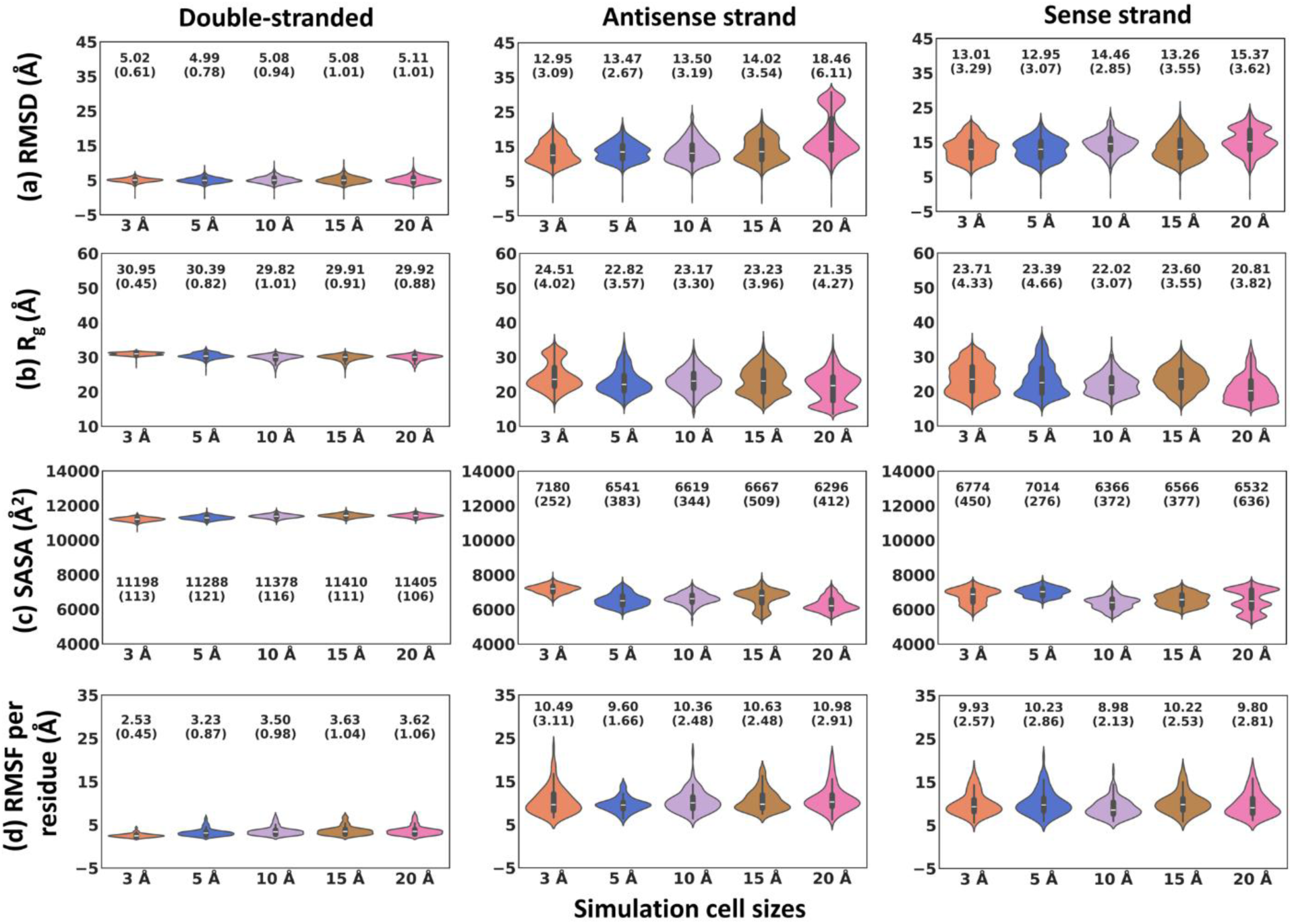
Violin plots showing the distribution of data across the full trajectory (1000 ns) for three DNA forms (double-stranded, antisense strand, sense strand) across five cell sizes (3 Å, 5 Å, 10 Å, 15 Å, 20 Å). Each violin represents 3,003 data points pooled from three independent MD replicates. Mean and standard deviation (in brackets) of these 3,003 data points are annotated with each violin. **(a)** Root-mean-square deviation (RMSD; in Å). **(b)** Radius of gyration (R_g_; in Å). **(c)** Solvent accessible surface area (SASA; in Å^2^). **(d)** Root-mean-square fluctuation per residue (RMSF; in Å).

A stronger cell-size effect was observed for the single-stranded systems. The largest average RMSD values were observed in the 20 Å cell (**Figure 2a**). This increase appears to be influenced by larger deviations in one replicate for each of the single-stranded systems, as reflected by the two distinct distributions in the violin plot for the 20 Å cell. In contrast, for the other cell sizes, the three replicates sampled more similar RMSD distributions, suggesting that these smaller cells restricted access to distinct conformational states. This indicates that the 20 Å may be more suitable for broader conformational sampling of single-stranded nucleic acids.

For the antisense strand (monomer 1), the average RMSD values for the 3, 5, 10, 15 and 20 Å cells were 12.95 ± 3.09, 13.47 ± 2.67, 13.50 ± 3.19, 14.02 ± 3.54, and 18.46 ± 6.11 Å, respectively. The corresponding values of the sense strand (monomer 2) were 13.01 ± 3.29, 12.95 ± 3.07, 14.46 ± 2.85, 13.26 ± 3.55, and 15.37 ± 3.62 Å (**Figures 2a** and **S1a**). Notably, higher RMSD values were observed in the 20 Å cell, while the smaller 3 and 5 Å cells generally showed lower RMSD values. This trend supports the interpretation that larger solvent buffers allow broader conformational sampling, while smaller cells restrict molecular motion. A similar but weaker trend was observed for the double-stranded system.

Comparison between the double-stranded and single-stranded systems showed substantially higher trajectory variability for the single-stranded molecules. The average RMSD values for the double-stranded DNA remained close to 5 Å, whereas those for the single-stranded systems ranged from approximately 13 to 18 Å. This suggests that the double-stranded DNA maintains greater structural stability than the isolated single strands. This behavior is expected because Watson-Crick base pairing and base stacking stabilize the double helix, while these stabilizing constraints are absent in the isolated single-stranded systems. As a result, the negatively charged phosphate backbone and exposed bases of the single strands interact more extensively with water and ions, leading to larger structural rearrangements. These observations also support the qualitative consistency of the protocol as the experimentally known facts can be clearly seen from double versus single stranded comparisons.

These analyses conclude that smaller cells (3 and 5 Å) restrict nucleic acid motion in both double-stranded and single-stranded systems. In contrast, larger cells offer more solvent space and allow broader conformational sampling. The effect is more pronounced for single-stranded nucleic acids than for the more stable double-helical system. These findings suggest that solvent-buffer distances larger than the commonly used 10 Å may be beneficial for sampling the conformational dynamics of flexible nucleic acids, especially single-stranded systems.

### 3.3 Single-stranded nucleic acids adopt compact conformations for stabilization

Consistent with the RMSD analysis, the average radius of gyration (R_g_) of the double-stranded DNA showed restricted variation in the 3 Å cell, while larger cells showed greater fluctuations (**Figures 2b** and **S1b**). The lowest standard deviation was observed for the 3 Å cell (0.45 Å), whereas the corresponding values for the other cell sizes ranged from 0.82 to 1.01 Å. However, the average R_g_ values were similar across all five cell sizes, ranging from 29.82 to 30.95 Å. This indicates that the overall compactness of the double-stranded DNA was largely maintained across the different solvent-buffer distances.

In contrast, the single-stranded systems showed lower R_g_ values (20.81 to 24.51 Å), particularly in the 20 Å cells. The lowest average R_g_ values were observed in the 20 Å cell, with values of 21.35 Å for the antisense strand and 20.81 Å for the sense strand. These lower R_g_ values suggest that the single-stranded DNA molecules adopt more compact conformations in larger cells, likely as a consequence of the loss of complementary base-pairing interactions. Such compaction may help stabilize the isolated strands by allowing intramolecular interactions, including base stacking, transient self-association, and water- or ion-mediated contacts.

This compaction tendency was observed in the single-stranded systems relative to the corresponding double-stranded systems across all cell sizes. However, the extent of compaction was more pronounced in the larger cells, where the increased solvent-buffer distance allows greater conformational freedom.

### 3.4 Larger cells support compact conformations of single-stranded nucleic acids

To understand the solvent exposure of the different nucleic acid systems, the solvent-accessible surface area (SASA) was calculated. For the double-helical structure, the average solvent exposure showed a slight increase with increasing cell size, although the differences were not pronounced. The SASA values for the 3, 5, 10, 15, and 20 Å cells were 11198 ± 113, 11288 ± 121, 11378 ± 116, 11410 ± 111, and 11405 ± 106 Å^2^, respectively. The highest average SASA was found for the 15 Å cell, although similar values were also obtained for the 10 and 20 Å cells. These results indicate that the double-stranded DNA maintained comparable solvent exposure across the different cell sizes, consistent with the structural stability of the duplex.

In contrast, the single-stranded DNA systems showed a notable decrease in SASA in the larger cells compared with the smaller cells (**Figures 2c** and **S2a**). For the antisense strand, the SASA values for the 3, 5, 10, 15 and 20 Å cells were 7180 ± 252, 6541 ± 383, 6619 ± 344, 6667 ± 509, and 6296 ± 412 Å^2^, respectively. The corresponding values for the sense strand were 6774 ± 450, 7014 ± 276, 6366 ± 372, 6566 ± 377, and 6532 ± 636 Å^2^. In the antisense strand, the lowest solvent exposure was observed in the 20 Å cell, whereas in case of sense strand, SASA remained broadly similar across the 10, 15 and 20 Å cells.

These results suggest that larger solvent-buffer distances allow single-stranded DNA to adopt more compact conformations, as reflected by the reduced SASA. This trend is consistent with the R_g_ analysis, where single-stranded systems showed lower R_g_ values in larger cells. The reduced SASA may arise from intramolecular compaction, transient self-association, base stacking, and water- or ion-mediated interactions that partially compensate for the loss of complementary base-pairing interactions. In the double-stranded DNA, cell-size-dependent effects were milder because the stable double-helical conformation restricts large structural rearrangements.

The polar surface area (PSA) followed a trend similar to SASA (see supporting information, **Figures S2b** and **S3a**). PSA remained relatively similar for the double-stranded systems, with only a minor increase in larger cells. In contrast, PSA decreased in the larger cells for the single-stranded systems, further supporting the formation of more compact conformations under extended solvent-buffer conditions.

### 3.5 R_g_ correlates with SASA in single-stranded molecule and in larger cells

R_g_ and SASA are known to correlate during protein simulations.[2] To examine whether a similar relationship is observed in nucleic acids, Pearson correlation coefficients (R-values) and p-values were calculated between frame-wise R_g_ and SASA values over the complete trajectories (**Figure S4**). Stronger meaningful correlations were observed for the single-stranded nucleic acids (R-value up to 0.92), whereas weak or negligible correlations were found for the double-stranded system (R-value up to 0.44). In the single-stranded systems, SASA increased with increasing R_g_, indicating that more expanded conformations expose a larger surface area to the solvent. In general, these correlations were stronger in the larger cells than in the smaller 3 and 5 Å cells. In contrast, the comparatively stable and structurally constrained double-stranded DNA showed limited variation in both R_g_ and SASA, resulting in weak or absent correlations. Similar trends were observed for the correlations between R_g_ and PSA (**Figure S5**).

These results suggest that the relationship between molecular compactness and solvent exposure is more clearly captured in single-stranded nucleic acids than in double-stranded DNA. This behavior is likely due to the greater conformational flexibility and compaction tendency of the single-stranded systems. The stronger correlations observed in larger cells further indicate that extended solvent-buffer distances allow single-stranded nucleic acids to sample a broader range of compact and extended conformations. Thus, larger cells may be more suitable for capturing the coupled size and surface-exposure dynamics of single-stranded nucleic acids.

### 3.6 Larger cells enhance nucleotide movements and dominant motions in duplex DNA

We further analyzed nucleotide-level movements across the five cell sizes using root-mean-square fluctuation (RMSF) over the full trajectories. As expected, terminal nucleotides showed higher fluctuations in all fifteen systems (**Figure S6**). In the double-stranded DNA systems, the smaller cells showed lower residue-level deviations, with average RMSF values of 2.53 and 3.23 Å in the 3 and 5 Å cells, respectively (**Figure 2d**). In contrast, the larger cells showed higher fluctuations, with average RMSF values of 3.50, 3.63, and 3.62 Å in the 10, 15, and 20 Å cells, respectively. However, the lower fluctuations in the smaller cells should not be interpreted as greater structural stability, as they likely arise from restricted solvent space and possible periodic image interactions that confine molecular motion.

In the single-stranded nucleic acid systems, significantly larger nucleotide movements were observed, with average RMSF values ranging from 8.98 to 10.98 Å (**Figure 2d**). This indicates greater flexibility of the isolated strands, which lack stabilizing base-pairing interactions and can undergo larger conformational rearrangements. These movements may also be associated with the transition of single-stranded molecules toward more compact conformations, as indicated by the R_g_ and SASA analyses.

To further characterize the dominant motions, free-energy landscape principal component analysis (FEL-PCA) was performed. In the double-helical systems, the 3 Å cell showed the smallest conformational landscape in the eigenvector 1 (EV1) versus eigenvector 2 (EV2) plots (**Figure S7**). This was also reflected in the lowest average EV1 and EV2 eigenvalues for the 3 Å cell, with values of 542 ± 228 and 461 ± 178 Å^2^, respectively (**Figure S8**). As the cell size increased, the EV1 and EV2 eigenvalues also increased, reaching maximum values of 1942 ± 305 and 1439 ± 37 Å^2^, respectively in the 20 Å cell. These results suggest that increasing the solvent-buffer distance enhances dominant collective motions in duplex DNA. In comparison, the single-stranded systems sampled broader conformational landscapes likely due to the absence of stabilizing base-pairing interactions (**Figures S7** and **S8**).

Together, the RMSF and FEL-PCA analyses show that larger solvent-buffer distances allow greater nucleotide-level and collective motions.

### 3.7 Non-covalent interactions support compact nucleic acid conformations in larger cells

The number of intra-nucleic acid hydrogen bonds remained stable in the double helical structure across all cell sizes (average = 79.26-80.29) (**Figures 3a** and **S9a**). This bonding also includes Watson-Crick base-pairing between complementary bases. In contrast, the single-stranded systems showed significantly fewer intra-nucleic acid hydrogen bonds (average = 14.89-19.58) due to the absence of complementary pairing. In the antisense strand, the highest hydrogen bonding content was observed in the 20 Å cell (average = 19.58), whereas the lowest value was observed in the 10 Å cell (average = 14.92). Similarly, in the sense strand, the highest hydrogen-bond content was observed in the 20 Å cell (average = 18.97), while the lowest value was found in the 5 Å cell (average = 14.89).

**FIGURE 3.**
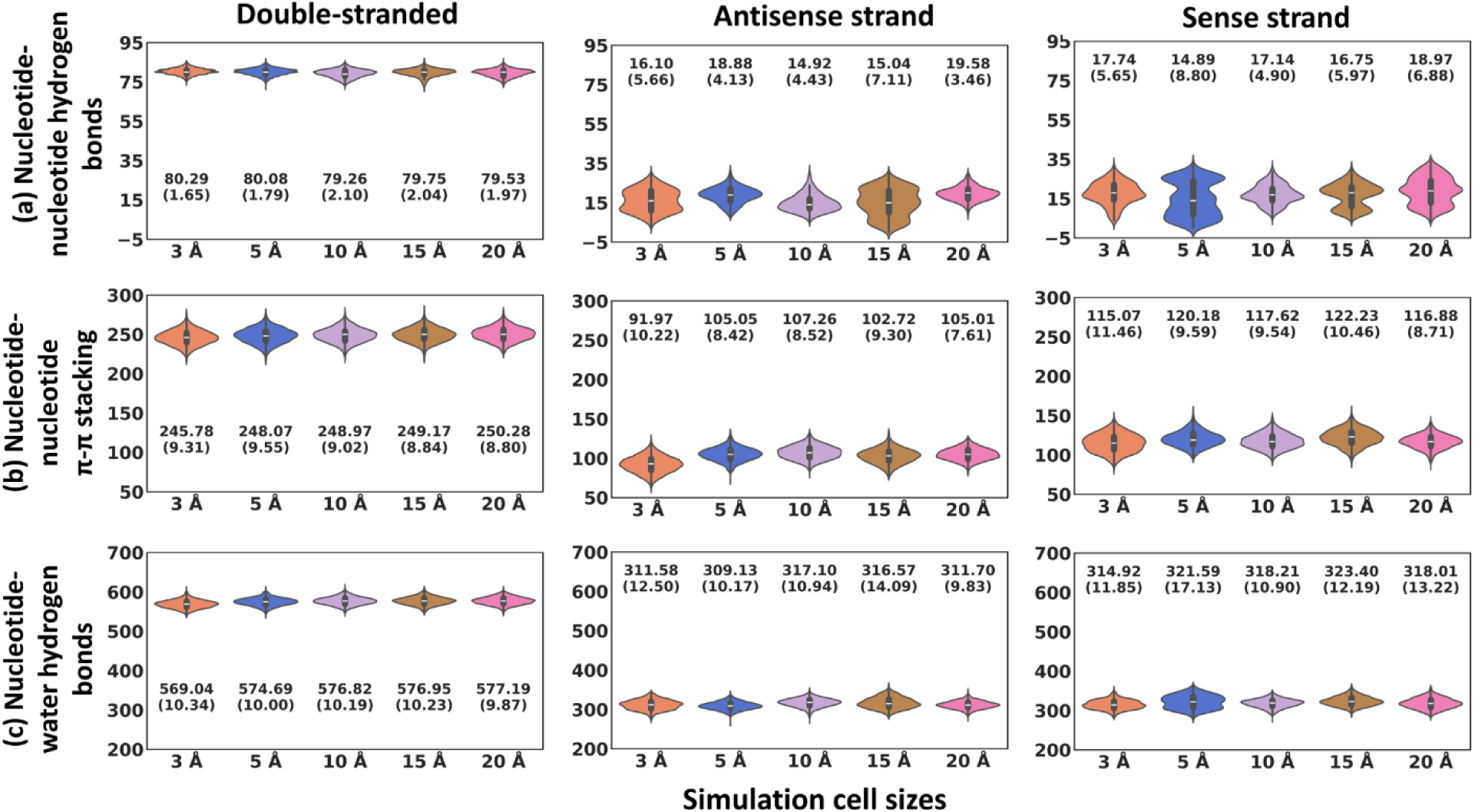
Violin plots showing the distribution of data across the equilibrated trajectory (last 500 ns) for three DNA forms (double-stranded, antisense strand, and sense strand) and five simulation cell sizes (3, 5, 10, 15, and 20 Å). Each violin represents 1,503 data points pooled from three independent MD replicates. Mean and standard deviation (in brackets) of these 1,503 data points are annotated with each violin. **(a)** Intramolecular nucleotide-nucleotide hydrogen bond count. **(b)** Nucleotide-nucleotide π-π stacking count. **(c)** Nucleotide-water hydrogen bond count.

These observations suggest that larger cells, particularly the 20 Å cell, allow single-stranded nucleic acids to form more intramolecular hydrogen bonds. This is consistent with the compact conformations observed from the R_g_ and SASA analyses, where larger solvent-buffer distances allowed the isolated strands to undergo self-association.

Consistent with the hydrogen bonding, π-π stacking interactions among nucleotides remained similar in the double-helical systems across the five cell sizes (average number = 245.78-250.28) (**Figures 3b** and **S9b**). The highest value was observed in the 20 Å cell (average = 250.28). Since π-π stacking between bases contributes to helix stabilization, the similar values across cell sizes further indicate that the double-helical structure remains comparatively stable. In the single-stranded systems, π-π stacking were more cell-size dependent. The antisense strand showed higher π-π interactions in the 10 and 20 Å cells, with average of 107.26 and 105.01, respectively. The sense strand showed the highest value in the 15 Å cell, with average of 122.23, although the 5 Å cell also showed a comparable value of 115.07. These results suggest that base-stacking interactions in single-stranded nucleic acids are more sensitive to solvent-buffer distance than those in the double helix.

The number of nucleic acid-water hydrogen bonds in the double-helical structure ranged from 569.04-577.19, with the lowest value observed in the 3 Å cell and the highest content in the 20 Å cell (**Figures 3c** and **S9c**). However, the differences among cell sizes were minor. In the single-stranded systems, this bonding ranged from 309.13 to 323.40. For the antisense strand, the 10 and 15 Å cells revealed higher water interactions, with average values of 317.10 and 316.57, respectively. For the sense strand, the highest value was observed in the 15 Å cell (average = 323.40). Interestingly, the 20 Å cells showed comparatively reduced water interactions in both single strands, with values of 311.70 and 318.01 for antisense and sense strands, respectively.

All these interactions conclude that larger cells (∼20 Å) provide sufficient solvent space for single-stranded nucleic acid to acquire a compact state. In the 20 Å cell, the increase in intra-nucleic acid hydrogen bonding and the reduction in nucleic acid-water hydrogen bonding are consistent with a more compact single-stranded conformation. These results support the use of larger solvent-buffer distances for studying flexible single-stranded nucleic acids, whereas the double-helical system is less sensitive to cell-size variation because of its stable base-paired and base-stacked structure.

### 3.8 Double-helical nucleic acid displays elongated conformation with variable strand lengths during simulations

To further investigate the dominant conformations, affinity propagation clustering was performed on the final 500 ns trajectories using a 3 Å RMSD cutoff for backbone atoms. The representative structures showed that the double-stranded DNA maintained an elongated rod-like conformation across all five cell sizes (**Figure 4**). In contrast, the single-stranded systems sampled transient compact and self-associated conformations in multiple simulation replicates, with the 20 Å cells showing a greater tendency toward such compact states.

**FIGURE 4.**
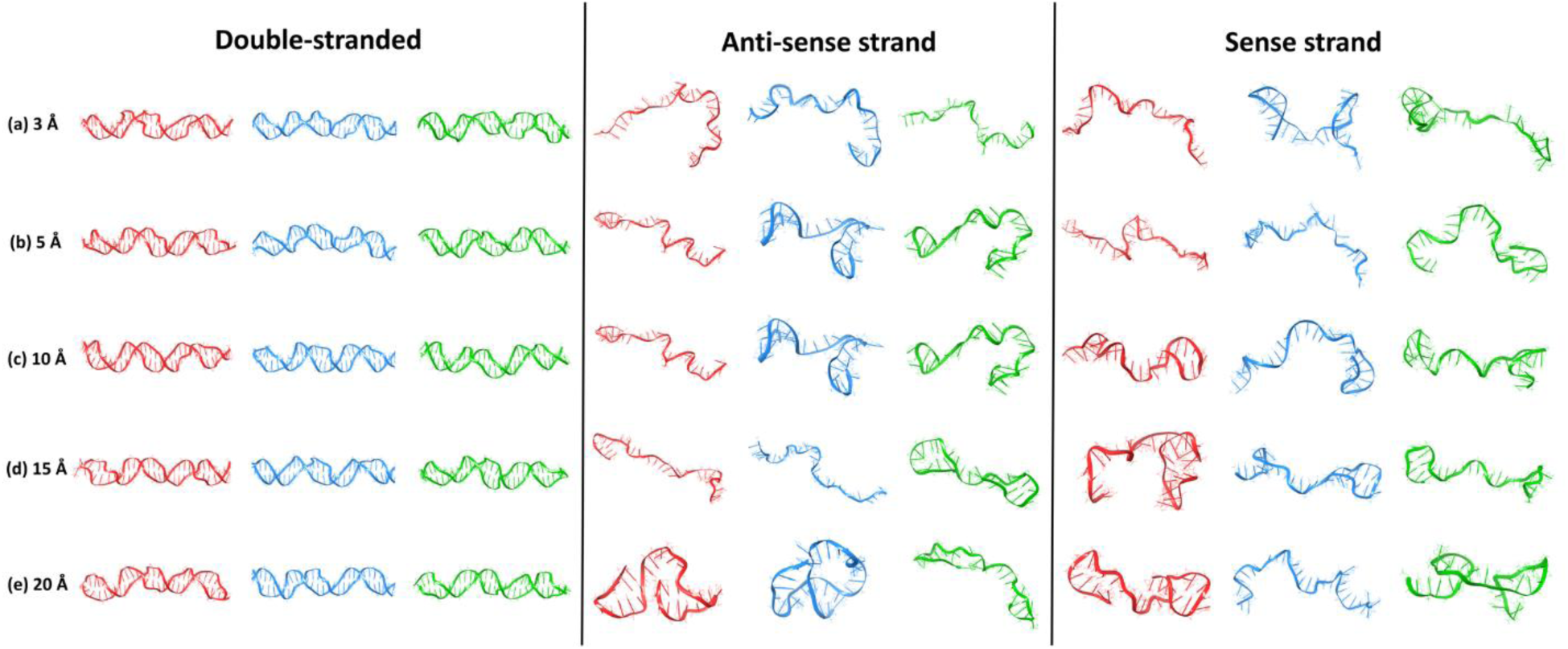
Representative structures from three independent MD replicates (MD1, MD2, MD3, shown in red, blue, and green, respectively) for the three DNA forms (double-stranded, antisense strand, and sense strand) at five simulation cell sizes: **(a)** 3 Å, **(b)** 5 Å, **(c)** 10 Å, **(d)** 15 Å, and **(e)** 20 Å.

To examine changes in duplex geometry, the apparent helical pitch was calculated as the distance between the first and tenth nucleotides of each strand. Because each strand contains 30 nucleotides, three pitch-like distances were calculated per strand, giving 18 measurements per cell size across three replicate simulations (3 pitches × 2 strands × 3 replicates) (**Figures S10**, **S11** and **Table S3**). In the experimental starting structure, the six pitch-like distances were similar, ranging from 32.29 to 34.11 Å. During simulations, these distances deviated from the starting structure and ranged from approximately 26 to 40 Å across the five cell sizes, indicating changes in duplex geometry. In the smallest 3 Å cell, the average first pitch-like distance decreased to 30.70 and 29.35 Å for antisense and sense strands, respectively (**Figure S11**). In contrast, the second pitch-like distance increased to 35.67 and 38.19 Å, and the third pitch-like distance increased to 37.87 and 34.45 Å for the antisense and sense strands, respectively.

These results were supported by terminal end-to-end distance analysis. The distances between the terminal nucleotide atoms, measured from the 5’ adenine to 3’ guanine in the antisense strand and from 5’ cytosine to 3’ thymine in the sense strand, were 101.90 and 102.36 Å, respectively, in the experimental structure (**Figures S12**, **S13** and **Table S7**). This indicates that both strands had similar end-to-end distances in the starting structure. During simulations, one strand in the double-helical system consistently showed a shorter end-to-end distance than the other, with values ranging from 94 to 109 Å. In contrast, all single-stranded simulations showed smaller terminal end-to-end distances, ranging from 16 to 100 Å, compared with the experimental end-to-end distance of approximately 102 Å. This supports the formation of compact conformations in the isolated single-stranded systems.

The helix width of the double-stranded DNA was further analyzed by calculating distances between atoms of adjacent nucleotides in complementary strands (**Figures S14**, **S15 and Table S4**). The average helix width remained close to the experimental structure across all cell sizes, with an average value of approximately 20 ± 0.25 Å. This indicates that although the duplex undergoes variations in pitch-like distances and terminal end-to-end distances during simulations, its overall interstrand width is largely maintained.

These observations indicate that the double-helical DNA retains an elongated duplex architecture, with one strand shorter than the other. In contrast, the single-stranded systems sample compact conformations, especially in the 20 Å cell. These results suggest that the commonly used 10 Å buffer may be sufficient for maintaining the overall double-helical architecture, whereas larger solvent-buffer distances are more suitable for sampling compact conformations of flexible single-stranded nucleic acids.

### 3.9 Nucleic acid shows reduced first-shell water density in larger cells

To examine the distribution of water molecules around the nucleic acid, radial distribution functions (RDFs) were calculated over the final 500 ns trajectories. RDF describes the probability of finding atoms or molecules at a given distance from a reference atom relative to the bulk density. Here, RDFs were calculated between the phosphorus atoms of DNA and the oxygen atoms of water, with the radial distance extended up to approximately half of the edge length *a* of the simulation cell (**Table S1**).

For both double-stranded and single-stranded systems, a sharp first peak was observed at approximately 3.8 Å radial distance, followed by a minimum and a smaller second peak at approximately 6 Å (**Figure 5a**). The first and second peaks correspond to the first and second hydration shells, respectively, while the intervening minimum defines the approximate boundary between these shells. Beyond these ordered shells, the RDF approached the bulk-water region.

**FIGURE 5.**
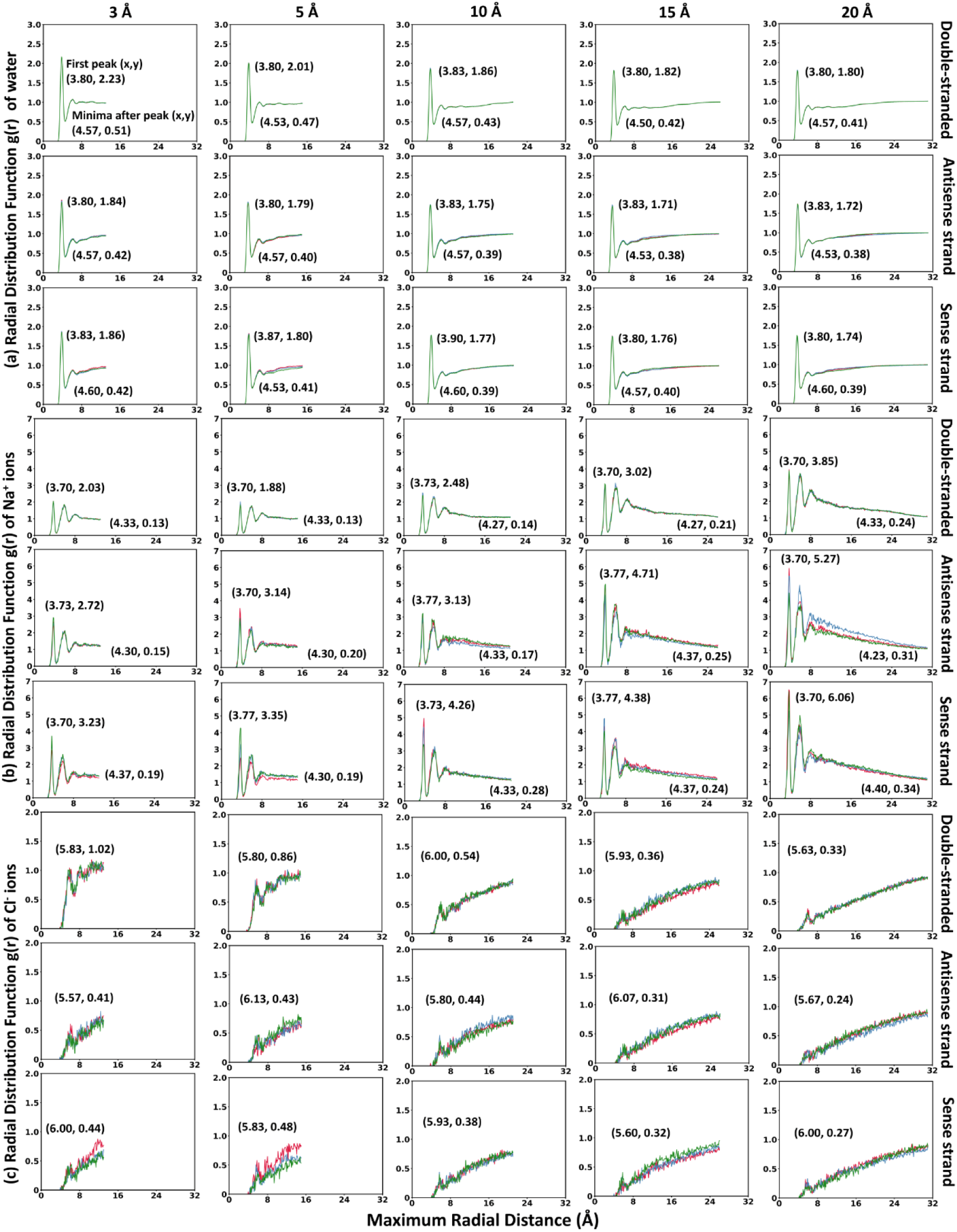
Radial distribution functions (RDF) of: **(a)** water molecules, **(b)** sodium ions, and **(c)** chloride ions around duplex DNA, and isolated antisense, and sense strands across five cell sizes (3, 5, 10, 15, and 20 Å). Three replicates are shown in red, blue, and green. The mean coordinates of the first solvation shell peak (x, y) and the subsequent minima (x, y), averaged across three replicates, are shown in each plot.

In the double-helical systems, the 3 Å cell showed a first-shell RDF peak of 2.23. This peak gradually decreased with increasing cell size and dropped to 1.80 in the 20 Å cell. Similar effects were observed for the isolated single strands. In the single-stranded systems, the first RDF peak remained below 1.9 across all cell sizes and decreased with increasing cell size (**Figure 5a**). For the antisense strand, the RDF peak decreased from 1.84 in the 3 Å cell to ∼1.71 in the 15 and 20 Å cells. Similarly, for the sense strand, the RDF peak decreased from 1.86 in the 3 Å cell to 1.74 in the 20 Å cell.

These findings suggest that smaller cells maintain a denser first hydration shell around the nucleic acid, likely due to restricted solvent space and confinement effects. In contrast, larger cells show reduced first-shell water density despite containing a larger total number of water molecules. This reduction may reflect lower molecular crowding around the DNA and greater conformational relaxation in larger solvent environments.

### 3.10 Cl^-^ ions show indirect association with nucleic acid-bound Na^+^ ions

To inspect ion distributions around the nucleic acid, RDFs were calculated between the phosphorus atoms of DNA and Na⁺ or Cl⁻ ions. Each system contained 0.15 M NaCl (**Table S2**); therefore, smaller cells contained fewer total ions than larger cells, although the ion concentration was kept constant.

For Na^+^ ions, three coordination shells were observed at approximately 3.7, 6.0 and 8.0 Å from the DNA phosphorus atoms (**Figure 5b**). As the cell size increased, the Na^+^ RDF peaks became more pronounced, indicating increased localization of Na^+^ ions around the phosphate groups. For the duplex DNA, the first RDF peak increased from 2.03 and 1.88 in 3 and 5 Å cells, respectively to 3.85 in the 20 Å cell. A similar trend was observed for the isolated strands. For the antisense strand, the first RDF peak increased from 2.72 in the 3 Å cell to 5.27 in the 20 Å cell.

For the sense strand, the corresponding peak increased from 3.23 in the 3 Å cell to 6.06 in the 20 Å cell. In all systems, the RDF values decreased at larger radial distances, showing that Na^+^ ions were preferentially distributed near the negatively charged phosphate backbone rather than in the bulk region. These observations are consistent with electrostatic association between Na^+^ ions and the phosphate groups of the nucleic acid backbone.

The distribution of Cl^-^ ions around nucleic acids showed a different trend (**Figure 5b**). A weak and distorted first-shell feature was observed at approximately 5.8 Å from the DNA phosphorus atoms. The smaller 3 and 5 Å cells showed higher apparent Cl^-^ RDF peaks, likely due to restricted ion mobility and confinement effects in the smaller solvent volume. With increasing cell size, the first-shell Cl^-^ feature became weaker and more diffuse. At larger radial distances, the Cl^-^ RDF increased toward the bulk region, indicating that Cl^-^ ions were less concentrated near the negatively charged nucleic acid backbone and more distributed away from the phosphate groups.

The weak Cl^-^ feature near 6 Å may reflect indirect association of Cl^-^ ions with Na^+^ ions localized near the phosphate backbone, rather than direct Cl^-^ binding to the nucleic acid. Thus, the RDF analysis suggests that Na^+^ ions preferentially associate with the phosphate groups, whereas Cl^-^ ions are largely excluded from the immediate phosphate environment and may appear near the nucleic acid through ion-pairing or solvent-mediated association with Na^+^ ions. In smaller cells, restricted solvent volume may artificially enhance local Cl^-^ density near the nucleic acid.

## 4 Conclusions

This study systematically investigated the effect of solvent-buffer distance on the molecular dynamics simulations of double-stranded and single-stranded DNA systems (**Figure 6**). For the double-helical DNA, the commonly used 10 Å solvent buffer was sufficient to maintain the overall duplex architecture, including stable hydrogen bonding, base stacking, and interstrand width. However, larger buffers allowed greater nucleotide-level fluctuations, broader collective motions, and partial helical relaxation, indicating improved conformational sampling.

**FIGURE 6.**
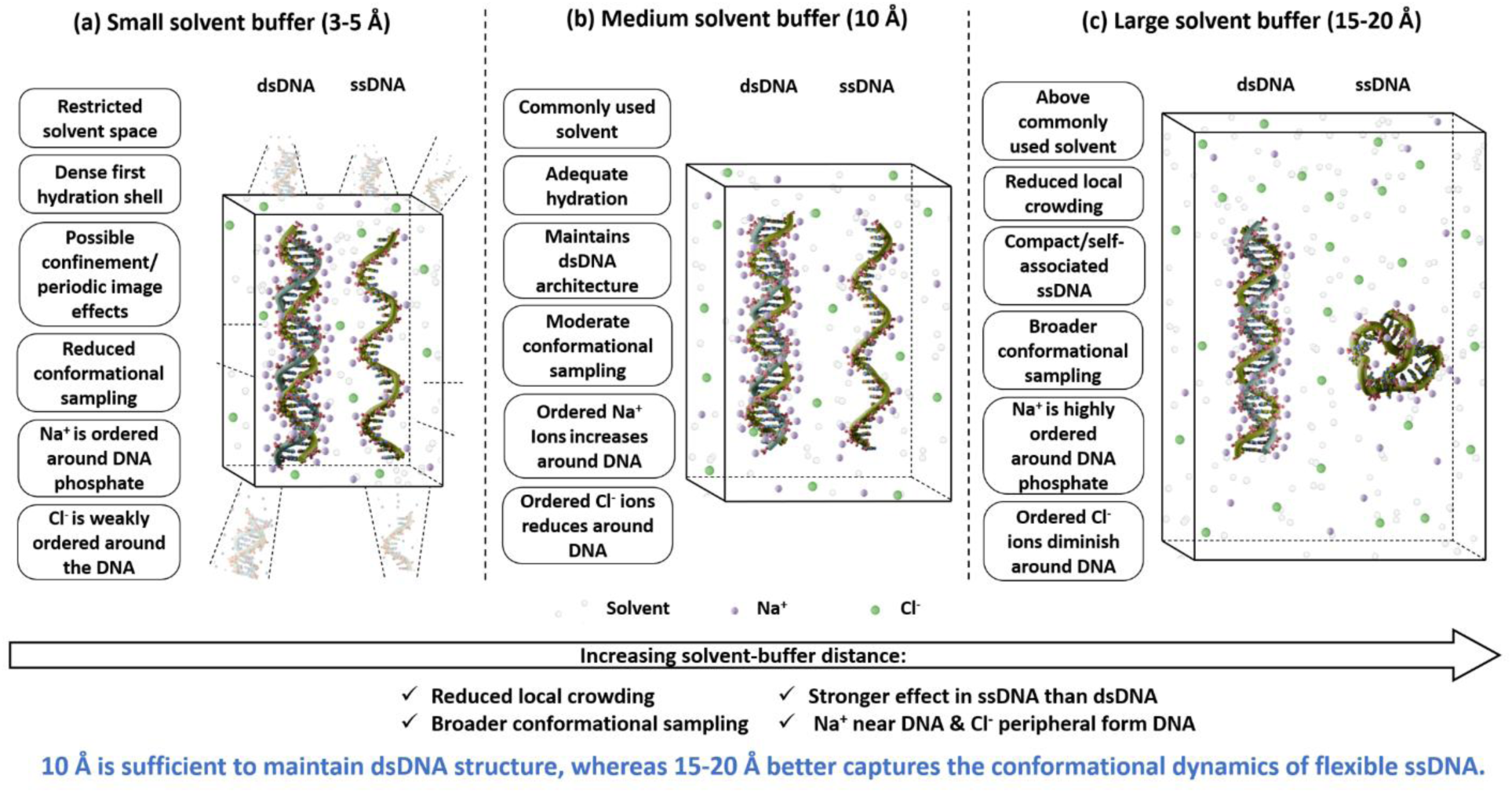
Summary of cell size effects in molecular dynamics simulations of nucleic acids. **(a)** Smaller solvent-buffer distances of 3-5 Å restrict conformational motion, increase local first-shell water density, and may enhance confinement or periodic image effects. **(b)** A 10 Å buffer is sufficient to maintain the overall architecture of double-stranded DNA. **(c)** Larger buffers of 15-20 Å allow broader conformational sampling, reduce local crowding, and are particularly important for flexible single-stranded DNA, which adopts compact and self-associated conformations. Ion distributions also vary with buffer size, with Na^+^ preferentially associating with the phosphate backbone and Cl^-^ largely excluded from the immediate phosphate environment. The double-stranded and single-stranded conformations are taken from the simulation representative structures.

The single-stranded DNA was more sensitive to cell-size variation. Larger solvent-buffer distances, particularly 15 and 20 Å, promote broader conformational sampling and compact, self-associated conformations. These compact states were supported by lower R_g_ and SASA values, stronger R_g_-SASA correlations, increased intramolecular hydrogen bonding, and changes in base-stacking and nucleic acid-water interactions. Thus, larger cells are more suitable for capturing the conformational dynamics of flexible single-stranded nucleic acids.

Hydration-shell and ion-distribution analyses further showed that smaller cells maintained a denser first hydration shell around the nucleic acid, likely due to restricted solvent space and confinement effects. With increasing cell size, the first-shell water density decreased, consistent with reduced local crowding and greater conformational relaxation. Na^+^ ions preferentially associated with the negatively charged phosphate backbone and showed ordered coordination shells around the nucleic acid, whereas Cl^-^ ions were largely excluded from the immediate phosphate environment and appeared to associate indirectly through Na^+^ ion-pairing or solvent-mediated interactions.

In brief, these results show that simulation cell size strongly influences nucleic acid dynamics, hydration structure, and ion organization. While a 10 Å solvent buffer may be adequate for maintaining the double-stranded DNA architecture, larger buffers are preferable for more comprehensively sampling the conformational dynamics of single-stranded nucleic acids. These findings provide practical guidance for choosing solvent-buffer distances in molecular dynamics simulations of nucleic acids.

The scope of this study should also be considered when interpreting the results. The present simulations were performed for DNA systems using the OPLS4 force field and the isolated, apo form of DNA without bound protein. RNA molecules may exhibit different cell-size dependence because of their distinct structural flexibility, base-pairing patterns, and hydration properties. Similarly, protein-nucleic acid complexes may behave differently because they combine protein structural constraints with the hydration and electrostatic features of nucleic acids. Different force fields may also produce somewhat different conformational and hydration behavior. Future studies on RNA systems, protein-nucleic acid complexes, and additional nucleic acid force fields will be useful for assessing the broader generality of the present observations.

## Acknowledgments

R.M. acknowledges Science and Engineering Research Board (ANRF/SERB), Government of India, for Start-up Research Grant SRG/2022/000304 and IIT Bhilai for Matching Grant, IITBhilai/D/2914. N.B. and P.S. acknowledge DST-INSPIRE for the fellowship.

## Author declarations

### Author Contribution

**Nainsy Baghel:** data curation, formal analysis, investigation, software, validation, visualization, writing review and editing. **Pranchal Shrivastava:** software, visualization, writing, reviewing, and editing. **Rukmankesh Mehra:** conceptualization, data curation, formal analysis, funding acquisition, investigation, methodology, project administration, resources, software, supervision, validation, visualization, writing original draft, and writing review and editing.

### Data Availability Statement

The data that support the findings of this study are available in the **Supporting Information** of this article.

The simulation data is available on GitHub:

https://github.com/Protein-informatics/cell_size_effects_in_nucleic_acids.git

