## Supplementary file for "Solvent-buffer effects in molecular dynamics simulations of nucleic acids"

*Nainsy Baghel:* 0009-0009-6024-3008

*Pranchal Shrivastava:* 0009-0005-4544-0346

*Rukmankesh Mehra:* 0000-0001-6010-1514

**Table S1. Dimensions and volume of the cell across each MD replicate.**

| Systems | | Edge *a* length | Edge *b* length | Edge *c* length | Cell volume |
|---|---|---|---|---|---|
| **Dimer (Double-stranded)** | | | | | |
| **3 Å** | **MD1** | 29.05 Å | 27.01 Å | 103.37 Å | 81121 Å$^3$ |
| | **MD2** | 29.04 Å | 27.00 Å | 103.32 Å | 81010 Å$^3$ |
| | **MD3** | 29.06 Å | 27.02 Å | 103.39 Å | 81168 Å$^3$ |
| **5 Å** | **MD1** | 33.53Å | 31.44 Å | 109.84 Å | 115797 Å$^3$ |
| | **MD2** | 33.56 Å | 31.47 Å | 109.94 Å | 116115 Å$^3$ |
| | **MD3** | 33.62 Å | 31.52 Å | 110.11 Å | 116674 Å$^3$ |
| **10 Å** | **MD1** | 44.35 Å | 42.19 Å | 123.45 Å | 231006 Å$^3$ |
| | **MD2** | 44.31Å | 42.15 Å | 123.33 Å | 230324 Å$^3$ |
| | **MD3** | 44.37 Å | 42.20 Å | 123.48 Å | 231171 Å$^3$ |
| **15 Å** | **MD1** | 54.49 Å | 52.30 Å | 134.37 Å | 382867 Å$^3$ |
| | **MD2** | 54.51 Å | 52.32 Å | 134.43 Å | 383404 Å$^3$ |
| | **MD3** | 54.48 Å | 52.29 Å | 134.35Å | 382685 Å$^3$ |
| **20 Å** | **MD1** | 64.74 Å | 62.53 Å | 145.32 Å | 588187 Å$^3$ |
| | **MD2** | 64.62 Å | 62.42 Å | 145.06 Å | 585074 Å$^3$ |
| | **MD3** | 64.71 Å | 62.50 Å | 145.26 Å | 587463 Å$^3$ |
| **Monomer 1 (Antisense strand)** | | | | | |
| **3 Å** | **MD1** | 27.21 Å | 28.82 Å | 103.73 Å | 81334 Å$^3$ |
| | **MD2** | 27.16 Å | 28.77 Å | 103.55 Å | 80924 Å$^3$ |
| | **MD3** | 27.20 Å | 28.81 Å | 103.68 Å | 81217 Å$^3$ |
| **5 Å** | **MD1** | 31.49 Å | 33.13 Å | 109.39 Å | 114102 Å$^3$ |
| | **MD2** | 31.63 Å | 33.28 Å | 109.87 Å | 115636 Å$^3$ |
| | **MD3** | 31.60 Å | 33.24 Å | 109.77 Å | 115307 Å$^3$ |
| **10 Å** | **MD1** | 42.03 Å | 43.71 Å | 121.96 Å | 224058 Å$^3$ |
| | **MD2** | 42.08 Å | 43.76 Å | 122.10 Å | 224851 Å$^3$ |
| | **MD3** | 42.11 Å | 43.79 Å | 122.19 Å | 225342 Å$^3$ |
| **15 Å** | **MD1** | 52.26 Å | 53.96 Å | 132.98 Å | 374979 Å$^3$ |
| | **MD2** | 52.20 Å | 53.89 Å | 132.82 Å | 373608 Å$^3$ |
| | **MD3** | 52.19 Å | 53.89 Å | 132.81 Å | 373574 Å$^3$ |
| **20 Å** | **MD1** | 62.16 Å | 63.87 Å | 143.00 Å | 567726 Å$^3$ |
| | **MD2** | 62.24 Å | 63.94 Å | 143.17 Å | 569682 Å$^3$ |
| | **MD3** | 64.98 Å | 62.83 Å | 143.07 Å | 584128 Å$^3$ |

| Monomer 2 (Sense strand) | | | | | |
|---|---|---|---|---|---|
| **3 Å** | **MD1** | 28.01 Å | 28.68 Å | 104.62 Å | 84031 $Å^3$ |
| | **MD2** | 28.02 Å | 28.69 Å | 104.67 Å | 84141 $Å^3$ |
| | **MD3** | 28.04 Å | 28.71 Å | 104.74 Å | 84312 $Å^3$ |
| **5 Å** | **MD1** | 32.11 Å | 32.78 Å | 109.59 Å | 115364 $Å^3$ |
| | **MD2** | 32.16 Å | 32.83 Å | 109.75 Å | 115865 $Å^3$ |
| | **MD3** | 32.22 Å | 32.90 Å | 109.97 Å | 116557 $Å^3$ |
| **10 Å** | **MD1** | 42.80 Å | 43.50 Å | 122.58 Å | 228205 $Å^3$ |
| | **MD2** | 42.78 Å | 43.48 Å | 122.53 Å | 227923 $Å^3$ |
| | **MD3** | 42.78 Å | 43.47 Å | 122.51 Å | 227795 $Å^3$ |
| **15 Å** | **MD1** | 52.79 Å | 53.49 Å | 132.96 Å | 375454 $Å^3$ |
| | **MD2** | 52.81 Å | 53.50 Å | 132.99 Å | 375740 $Å^3$ |
| | **MD3** | 52.77 Å | 53.47 Å | 132.90 Å | 375011 $Å^3$ |
| **20 Å** | **MD1** | 62.71 Å | 63.41 Å | 143.04 Å | 568775 $Å^3$ |
| | **MD2** | 62.82 Å | 63.52 Å | 143.30 Å | 571884 $Å^3$ |
| | **MD3** | 62.79 Å | 63.49 Å | 143.22 Å | 570902 $Å^3$ |

**Table S2. The number of atoms and residues in each MD replicate.**

| Systems | | All | | DNA | | Solvent | | Salt ions |
|---|---|---|---|---|---|---|---|---|
| | | Atom | Residue | Atom | Residue | Atom | Residue | Atom |
| 3 Å | MD1 | 8360 | 2260 | 1904 | 60 | 6384 | 2128 | 72 |
| | MD2 | 8360 | 2260 | 1904 | 60 | 6384 | 2128 | 72 |
| | MD3 | 8360 | 2260 | 1904 | 60 | 6384 | 2128 | 72 |
| 5 Å | MD1 | 11807 | 3413 | 1904 | 60 | 9825 | 3275 | 78 |
| | MD2 | 11807 | 3413 | 1904 | 60 | 9825 | 3275 | 78 |
| | MD3 | 11807 | 3413 | 1904 | 60 | 9825 | 3275 | 78 |
| 10 Å | MD1 | 23124 | 7200 | 1904 | 60 | 21120 | 7040 | 100 |
| | MD2 | 23124 | 7200 | 1904 | 60 | 21120 | 7040 | 100 |
| | MD3 | 23124 | 7200 | 1904 | 60 | 21120 | 7040 | 100 |
| 15 Å | MD1 | 38132 | 12220 | 1904 | 60 | 36102 | 12034 | 126 |
| | MD2 | 38132 | 12220 | 1904 | 60 | 36102 | 12034 | 126 |
| | MD3 | 38132 | 12220 | 1904 | 60 | 36102 | 12034 | 126 |
| 20 Å | MD1 | 58216 | 18940 | 1904 | 60 | 56148 | 18716 | 164 |
| | MD2 | 58216 | 18940 | 1904 | 60 | 56148 | 18716 | 164 |
| | MD3 | 58216 | 18940 | 1904 | 60 | 56148 | 18716 | 164 |
| **Monomer 1 (Antisense strand)** | | | | | | | | |
| 3 Å | MD1 | 8175 | 2469 | 946 | 30 | 7185 | 2395 | 44 |
| | MD2 | 8175 | 2469 | 946 | 30 | 7185 | 2395 | 44 |
| | MD3 | 8175 | 2469 | 946 | 30 | 7185 | 2395 | 44 |
| 5 Å | MD1 | 11541 | 3595 | 946 | 30 | 10545 | 3515 | 50 |
| | MD2 | 11541 | 3595 | 946 | 30 | 10545 | 3515 | 50 |
| | MD3 | 11541 | 3595 | 946 | 30 | 10545 | 3515 | 50 |
| 10 Å | MD1 | 22379 | 7221 | 946 | 30 | 21363 | 7121 | 70 |
| | MD2 | 22379 | 7221 | 946 | 30 | 21363 | 7121 | 70 |
| | MD3 | 22379 | 7221 | 946 | 30 | 21363 | 7121 | 70 |
| 15 Å | MD1 | 37065 | 12135 | 946 | 30 | 36021 | 12007 | 98 |
| | MD2 | 37065 | 12135 | 946 | 30 | 36021 | 12007 | 98 |

| | | | | | | | | |
|---|---|---|---|---|---|---|---|---|
| | **MD3** | 37065 | 12135 | 946 | 30 | 36021 | 12007 | 98 |
| **20 Å** | **MD1** | 56239 | 18549 | 946 | 30 | 55161 | 18387 | 132 |
| | **MD2** | 56239 | 18549 | 946 | 30 | 55161 | 18387 | 132 |
| | **MD3** | 57710 | 19042 | 946 | 30 | 56628 | 18876 | 136 |
| **3 Å** | **MD1** | 8496 | 2572 | 958 | 30 | 7494 | 2498 | 44 |
| | **MD2** | 8496 | 2572 | 958 | 30 | 7494 | 2498 | 44 |
| | **MD3** | 8496 | 2572 | 958 | 30 | 7494 | 2498 | 44 |
| **5 Å** | **MD1** | 11658 | 3630 | 958 | 30 | 10650 | 3550 | 50 |
| | **MD2** | 11658 | 3630 | 958 | 30 | 10650 | 3550 | 50 |
| | **MD3** | 11658 | 3630 | 958 | 30 | 10650 | 3550 | 50 |
| **10 Å** | **MD1** | 22649 | 7307 | 958 | 30 | 21621 | 7207 | 70 |
| | **MD2** | 22649 | 7307 | 958 | 30 | 21621 | 7207 | 70 |
| | **MD3** | 22649 | 7307 | 958 | 30 | 21621 | 7207 | 70 |
| **15 Å** | **MD1** | 37218 | 12182 | 958 | 30 | 36162 | 12054 | 98 |
| | **MD2** | 37218 | 12182 | 958 | 30 | 36162 | 12054 | 98 |
| | **MD3** | 37218 | 12182 | 958 | 30 | 36162 | 12054 | 98 |
| **20 Å** | **MD1** | 56293 | 18563 | 958 | 30 | 55023 | 18563 | 132 |
| | **MD2** | 56293 | 18563 | 958 | 30 | 55023 | 18563 | 132 |
| | **MD3** | 56293 | 18563 | 958 | 30 | 55023 | 18563 | 132 |

**Table S3. The helical pitch (Å) of each double-stranded DNA measured from representative structure of each MD replicate.**

| Systems | | | | Distance |
|---|---|---|---|---|
| **PDB ID: 7EDS** | | Monomer 1 | 1st Adenine to 10th Thymine | 32.40 Å |
| | | | 10th Thymine to 20th Adenine | 33.94 Å |
| | | | 20th Adenine to 30th Guanine | 33.73 Å |
| | | Monomer 2 | 1st Cytosine to 10th Adenine | 32.29 Å |
| | | | 10thAdenine to 20thAdenine | 34.11 Å |
| | | | 20th Adenine to 30th Thymine | 34.02 Å |
| | **Dimer (Double-stranded)** | | | |
| **3 Å** | **MD1** | Monomer 1 | 1st Adenine to 10th Thymine | 31.33 Å |
| | | | 10th Thymine to 20th Adenine | 35.21 Å |
| | | | 20th Adenine to 30th Guanine | 37.09 Å |
| | | Monomer 2 | 1st Cytosine to 10th Adenine | 29.65 Å |
| | | | 10thAdenine to 20thAdenine | 39.66 Å |
| | | | 20th Adenine to 30th Thymine | 35.95 Å |
| | **MD2** | Monomer 1 | 1st Adenine to 10th Thymine | 30.06 Å |
| | | | 10th Thymine to 20th Adenine | 36.98 Å |
| | | | 20th Adenine to 30th Guanine | 38.44 Å |
| | | Monomer 2 | 1st Cytosine to 10th Adenine | 28.98 Å |
| | | | 10thAdenine to 20thAdenine | 37.39 Å |
| | | | 20th Adenine to 30th Thymine | 36.59 Å |
| | **MD3** | Monomer 1 | 1st Adenine to 10th Thymine | 29. 84 Å |
| | | | 10th Thymine to 20th Adenine | 34.81 Å |
| | | | 20th Adenine to 30th Guanine | 38.07 Å |
| | | Monomer 2 | 1st Cytosine to 10th Adenine | 29.42 Å |
| | | | 10thAdenine to 20thAdenine | 37.52 Å |
| | | | 20th Adenine to 30th Thymine | 30.82 Å |
| **5 Å** | **MD1** | Monomer 1 | 1st Adenine to 10th Thymine | 25.83 Å |
| | | | 10th Thymine to 20th Adenine | 35.24 Å |
| | | | 20th Adenine to 30th Guanine | 37.62 Å |
| | | Monomer 2 | 1st Cytosine to 10th Adenine | 36.84 Å |
| | | | 10thAdenine to 20thAdenine | 37.11 Å |
| | | | 20th Adenine to 30th Thymine | 36.66 Å |
| | **MD2** | Monomer 1 | 1st Adenine to 10th Thymine | 33.13 Å |
| | | | 10th Thymine to 20th Adenine | 35.14 Å |
| | | | 20th Adenine to 30th Guanine | 33.21 Å |
| | | Monomer 2 | 1st Cytosine to 10th Adenine | 34.32 Å |
| | | | 10thAdenine to 20thAdenine | 37.01 Å |
| | | | 20th Adenine to 30th Thymine | 37.63 Å |

| | | | | |
|---|---|---|---|---|
| | MD3 | Monomer 1 | 1st Adenine to 10th Thymine | 36.56 Å |
| | | | 10th Thymine to 20th Adenine | 34.93 Å |
| | | | 20th Adenine to 30th Guanine | 36.65 Å |
| | | Monomer 2 | 1st Cytosine to 10th Adenine | 30.06 Å |
| | | | 10th Adenine to 20th Adenine | 35.99 Å |
| | | | 20th Adenine to 30th Thymine | 35.52 Å |
| **10 Å** | **MD1** | Monomer 1 | 1st Adenine to 10th Thymine | 33.41 Å |
| | | | 10th Thymine to 20th Adenine | 35.47 Å |
| | | | 20th Adenine to 30th Guanine | 36.62 Å |
| | | Monomer 2 | 1st Cytosine to 10th Adenine | 33.47 Å |
| | | | 10th Adenine to 20th Adenine | 34.07 Å |
| | | | 20th Adenine to 30th Thymine | 32.09 Å |
| | **MD2** | Monomer 1 | 1st Adenine to 10th Thymine | 34.96 Å |
| | | | 10th Thymine to 20th Adenine | 35.81 Å |
| | | | 20th Adenine to 30th Guanine | 33.36 Å |
| | | Monomer 2 | 1st Cytosine to 10th Adenine | 34.23 Å |
| | | | 10th Adenine to 20th Adenine | 31.70 Å |
| | | | 20th Adenine to 30th Thymine | 34.80 Å |
| | **MD3** | Monomer 1 | 1st Adenine to 10th Thymine | 30.78 Å |
| | | | 10th Thymine to 20th Adenine | 38.41 Å |
| | | | 20th Adenine to 30th Guanine | 37.41 Å |
| | | Monomer 2 | 1st Cytosine to 10th Adenine | 32.02 Å |
| | | | 10th Adenine to 20th Adenine | 31.24 Å |
| | | | 20th Adenine to 30th Thymine | 32.64 Å |
| **15 Å** | **MD1** | Monomer 1 | 1st Adenine to 10th Thymine | 34.80 Å |
| | | | 10th Thymine to 20th Adenine | 35.70 Å |
| | | | 20th Adenine to 30th Guanine | 35.41 Å |
| | | Monomer 2 | 1st Cytosine to 10th Adenine | 28.50 Å |
| | | | 10th Adenine to 20th Adenine | 34.22 Å |
| | | | 20th Adenine to 30th Thymine | 38.77 Å |
| | **MD2** | Monomer 1 | 1st Adenine to 10th Thymine | 36.25 Å |
| | | | 10th Thymine to 20th Adenine | 36.14 Å |
| | | | 20th Adenine to 30th Guanine | 35.60 Å |
| | | Monomer 2 | 1st Cytosine to 10th Adenine | 31.10 Å |
| | | | 10th Adenine to 20th Adenine | 37.87 Å |
| | | | 20th Adenine to 30th Thymine | 31.07 Å |
| | **MD3** | Monomer 1 | 1st Adenine to 10th Thymine | 28.39 Å |
| | | | 10th Thymine to 20th Adenine | 35.07 Å |
| | | | 20th Adenine to 30th Guanine | 38.17 Å |
| | | Monomer 2 | 1st Cytosine to 10th Adenine | 27.43 Å |
| | | | 10th Adenine to 20th Adenine | 35.31 Å |

| | | | | |
|---|---|---|---|---|
| | | | 20$^{th}$ Adenine to 30$^{th}$ Thymine | 34.61 Å |
| **20 Å** | **MD1** | Monomer 1 | 1$^{st}$ Adenine to 10$^{th}$ Thymine | 29.00 Å |
| | | | 10$^{th}$ Thymine to 20$^{th}$ Adenine | 34.06 Å |
| | | | 20$^{th}$ Adenine to 30$^{th}$ Guanine | 33.70 Å |
| | | Monomer 2 | 1$^{st}$ Cytosine to 10$^{th}$ Adenine | 37.20 Å |
| | | | 10$^{th}$Adenine to 20$^{th}$Adenine | 36.78 Å |
| | | | 20$^{th}$ Adenine to 30$^{th}$ Thymine | 38.97 Å |
| | **MD2** | Monomer 1 | 1$^{st}$ Adenine to 10$^{th}$ Thymine | 31.68 Å |
| | | | 10$^{th}$ Thymine to 20$^{th}$ Adenine | 35.82 Å |
| | | | 20$^{th}$ Adenine to 30$^{th}$ Guanine | 32.11 Å |
| | | Monomer 2 | 1$^{st}$ Cytosine to 10$^{th}$ Adenine | 35.06 Å |
| | | | 10$^{th}$Adenine to 20$^{th}$Adenine | 35.62 Å |
| | | | 20$^{th}$ Adenine to 30$^{th}$ Thymine | 31.00 Å |
| | **MD3** | Monomer 1 | 1$^{st}$ Adenine to 10$^{th}$ Thymine | 33.96 Å |
| | | | 10$^{th}$ Thymine to 20$^{th}$ Adenine | 34.27 Å |
| | | | 20$^{th}$ Adenine to 30$^{th}$ Guanine | 39.38 Å |
| | | Monomer 2 | 1$^{st}$ Cytosine to 10$^{th}$ Adenine | 28.16 Å |
| | | | 10$^{th}$Adenine to 20$^{th}$Adenine | 37.87 Å |
| | | | 20$^{th}$ Adenine to 30$^{th}$ Thymine | 36.58 Å |

**Table S4. The helical diameter (Å) of each double-stranded DNA measured from representative structure of each MD replicate.**

| Systems | | Selected atom: Monomer 1 (Antisense strand) | Selected atom: Monomer 2 (Sense strand) | distance |
|---|---|---|---|---|
| PDB ID: 7EDS | | $3^{rd}$ position Adenine | $28^{th}$ position Thymine | 19.8 Å |
| | | $8^{th}$ position Guanine | $23^{rd}$ position Cytosine | 20.03 Å |
| | | $13^{th}$ position Cytosine | $18^{th}$ position Guanine | 20.06 Å |
| | | $18^{th}$ position Cytosine | $13^{th}$ position Guanine | 19.68 Å |
| | | $23^{rd}$ position Cytosine | $8^{th}$ position Guanine | 20.36 Å |
| | | $28^{th}$ position Guanine | $3^{rd}$ position Cytosine | 20.21 Å |
| Dimer (Double-stranded) | | | | |
| 3 Å | MD1 | $3^{rd}$ position Adenine | $28^{th}$ position Thymine | 20.32 Å |
| | | $8^{th}$ position Guanine | $23^{rd}$ position Cytosine | 19.7 Å |
| | | $13^{th}$ position Cytosine | $18^{th}$ position Guanine | 19.53 Å |
| | | $18^{th}$ position Cytosine | $13^{th}$ position Guanine | 19.18 Å |
| | | $23^{rd}$ position Cytosine | $8^{th}$ position Guanine | 21.05 Å |
| | | $28^{th}$ position Guanine | $3^{rd}$ position Cytosine | 17.79 Å |
| | MD2 | $3^{rd}$ position Adenine | $28^{th}$ position Thymine | 19.42 Å |
| | | $8^{th}$ position Guanine | $23^{rd}$ position Cytosine | 19.53 Å |
| | | $13^{th}$ position Cytosine | $18^{th}$ position Guanine | 18.57 Å |
| | | $18^{th}$ position Cytosine | $13^{th}$ position Guanine | 18.44 Å |
| | | $23^{rd}$ position Cytosine | $8^{th}$ position Guanine | 19.11 Å |
| | | $28^{th}$ position Guanine | $3^{rd}$ position Cytosine | 19.8 Å |
| | MD3 | $3^{rd}$ position Adenine | $28^{th}$ position Thymine | 19.22 Å |
| | | $8^{th}$ position Guanine | $23^{rd}$ position Cytosine | 19.84 Å |
| | | $13^{th}$ position Cytosine | $18^{th}$ position Guanine | 20.79 Å |
| | | $18^{th}$ position Cytosine | $13^{th}$ position Guanine | 17.15 Å |
| | | $23^{rd}$ position Cytosine | $8^{th}$ position Guanine | 20.86 Å |
| | | $28^{th}$ position Guanine | $3^{rd}$ position Cytosine | 19.64 Å |
| 5 Å | MD1 | $3^{rd}$ position Adenine | $28^{th}$ position Thymine | 19.35 Å |
| | | $8^{th}$ position Guanine | $23^{rd}$ position Cytosine | 19.07 Å |
| | | $13^{th}$ position Cytosine | $18^{th}$ position Guanine | 20.81 Å |
| | | $18^{th}$ position Cytosine | $13^{th}$ position Guanine | 21.93 Å |
| | | $23^{rd}$ position Cytosine | $8^{th}$ position Guanine | 20.22 Å |
| | | $28^{th}$ position Guanine | $3^{rd}$ position Cytosine | 20.08 Å |
| | MD2 | $3^{rd}$ position Adenine | $28^{th}$ position Thymine | 18.33 Å |
| | | $8^{th}$ position Guanine | $23^{rd}$ position Cytosine | 19.98 Å |
| | | $13^{th}$ position Cytosine | $18^{th}$ position Guanine | 20.53 Å |
| | | $18^{th}$ position Cytosine | $13^{th}$ position Guanine | 20.96 Å |
| | | $23^{rd}$ position Cytosine | $8^{th}$ position Guanine | 18.81 Å |
| | | $28^{th}$ position Guanine | $3^{rd}$ position Cytosine | 19.15 Å |

| | | | | |
|---|---|---|---|---|
| | **MD3** | 3rd position Adenine | 28th position Thymine | 21.47 Å |
| | | 8th position Guanine | 23rd position Cytosine | 17.61 Å |
| | | 13th position Cytosine | 18th position Guanine | 21.52 Å |
| | | 18th position Cytosine | 13th position Guanine | 21.94 Å |
| | | 23rd position Cytosine | 8th position Guanine | 18.65 Å |
| | | 28th position Guanine | 3rd position Cytosine | 20.7 Å |
| **10 Å** | **MD1** | 3rd position Adenine | 28th position Thymine | 20.34 Å |
| | | 8th position Guanine | 23rd position Cytosine | 19.94 Å |
| | | 13th position Cytosine | 18th position Guanine | 18.51 Å |
| | | 18th position Cytosine | 13th position Guanine | 19.19 Å |
| | | 23rd position Cytosine | 8th position Guanine | 20.79 Å |
| | | 28th position Guanine | 3rd position Cytosine | 20.06 Å |
| | **MD2** | 3rd position Adenine | 28th position Thymine | 21.23 Å |
| | | 8th position Guanine | 23rd position Cytosine | 19.11 Å |
| | | 13th position Cytosine | 18th position Guanine | 19.63 Å |
| | | 18th position Cytosine | 13th position Guanine | 20.57 Å |
| | | 23rd position Cytosine | 8th position Guanine | 19.64 Å |
| | | 28th position Guanine | 3rd position Cytosine | 19.61 Å |
| | **MD3** | 3rd position Adenine | 28th position Thymine | 19.68 Å |
| | | 8th position Guanine | 23rd position Cytosine | 20.42 Å |
| | | 13th position Cytosine | 18th position Guanine | 19.47 Å |
| | | 18th position Cytosine | 13th position Guanine | 19.65 Å |
| | | 23rd position Cytosine | 8th position Guanine | 19.17 Å |
| | | 28th position Guanine | 3rd position Cytosine | 20.81 Å |
| **15 Å** | **MD1** | 3rd position Adenine | 28th position Thymine | 16.92 Å |
| | | 8th position Guanine | 23rd position Cytosine | 18.63 Å |
| | | 13th position Cytosine | 18th position Guanine | 19.59 Å |
| | | 18th position Cytosine | 13th position Guanine | 19.22 Å |
| | | 23rd position Cytosine | 8th position Guanine | 19.89 Å |
| | | 28th position Guanine | 3rd position Cytosine | 17.6 Å |
| | **MD2** | 3rd position Adenine | 28th position Thymine | 18.89 Å |
| | | 8th position Guanine | 23rd position Cytosine | 21.82 Å |
| | | 13th position Cytosine | 18th position Guanine | 20.91 Å |
| | | 18th position Cytosine | 13th position Guanine | 19.87 Å |
| | | 23rd position Cytosine | 8th position Guanine | 19.88 Å |
| | | 28th position Guanine | 3rd position Cytosine | 19.12 Å |
| | **MD3** | 3rd position Adenine | 28th position Thymine | 20.53 Å |
| | | 8th position Guanine | 23rd position Cytosine | 18.79 Å |
| | | 13th position Cytosine | 18th position Guanine | 19.28 Å |
| | | 18th position Cytosine | 13th position Guanine | 21.03 Å |
| | | 23rd position Cytosine | 8th position Guanine | 20.43 Å |
| | | 28th position Guanine | 3rd position Cytosine | 19.54 Å |
| **20 Å** | **MD1** | 3rd position Adenine | 28th position Thymine | 20.66 Å |

| | | | | |
|---|---|---|---|---|
| | | $8^{th}$ position Guanine | $23^{rd}$ position Cytosine | 20.79 Å |
| | | $13^{th}$ position Cytosine | $18^{th}$ position Guanine | 20.87 Å |
| | | $18^{th}$ position Cytosine | $13^{th}$ position Guanine | 19.78 Å |
| | | $23^{rd}$ position Cytosine | $8^{th}$ position Guanine | 20.36 Å |
| | | $28^{th}$ position Guanine | $3^{rd}$ position Cytosine | 20.55 Å |
| | **MD2** | $3^{rd}$ position Adenine | $28^{th}$ position Thymine | 20.82 Å |
| | | $8^{th}$ position Guanine | $23^{rd}$ position Cytosine | 18.13 Å |
| | | $13^{th}$ position Cytosine | $18^{th}$ position Guanine | 19.22 Å |
| | | $18^{th}$ position Cytosine | $13^{th}$ position Guanine | 20.36 Å |
| | | $23^{rd}$ position Cytosine | $8^{th}$ position Guanine | 19.36 Å |
| | | $28^{th}$ position Guanine | $3^{rd}$ position Cytosine | 20.29 Å |
| | **MD3** | $3^{rd}$ position Adenine | $28^{th}$ position Thymine | 19.04 Å |
| | | $8^{th}$ position Guanine | $23^{rd}$ position Cytosine | 19.48 Å |
| | | $13^{th}$ position Cytosine | $18^{th}$ position Guanine | 19.04 Å |
| | | $18^{th}$ position Cytosine | $13^{th}$ position Guanine | 21.19 Å |
| | | $23^{rd}$ position Cytosine | $8^{th}$ position Guanine | 20.75 Å |
| | | $28^{th}$ position Guanine | $3^{rd}$ position Cytosine | 20.09 Å |

**Table S5. Sequence analysis of all three initial DNA forms.**

| Parameter | Values |
|---|---|
| **Dimer (Double-stranded)** | |
| GC % | 66.67% |
| A | 16.67%, 10 residues |
| C | 33.33%, 20 residues |
| G | 33.33%, 20 residues |
| T | 16.67%, 10 residues |
| **Monomer 1 (Antisense strand)** | |
| GC % | 66.67% |
| A | 16.67%, 5 residues |
| C | 40%, 12 residues |
| G | 26.67% 8 residues |
| T | 16.67% 5 residues |
| **Monomer 2 (Sense strand)** | |
| GC % | 66.67% |
| A | 16.67%, 5 residues |
| C | 26.67%, 8 residues |
| G | 40% 12 residues |
| T | 16.67% 5 residues |

**Table S6. Analysis of variance (ANOVA test). A p-value ≤ 0.05 indicates statistically significant difference between the groups.**

| Dimer (Double-stranded) | | | |
|---|---|---|---|
| | **Group Pair** | **p-value** | **Group pair significance** |
| **RMSD** Total (1001*3) =3003 datapoints | **CellSize_3Å - CellSize_5Å** | **0.77** | **Statistically non-significant** |
| | CellSize_3Å - CellSize_10Å | 0.02 | Statistically significant |
| | **CellSize_3Å - CellSize_15Å** | **0.06** | **Statistically non-significant** |
| | CellSize_3Å - CellSize_20Å | 0.00 | Statistically significant |
| | CellSize_5Å - CellSize_10Å | 0.00 | Statistically significant |
| | CellSize_5Å - CellSize_15Å | 0.00 | Statistically significant |
| | CellSize_5Å - CellSize_20Å | 0.00 | Statistically significant |
| | **CellSize_10Å - CellSize_15Å** | **1.00** | **Statistically non-significant** |
| | **CellSize_10Å - CellSize_20Å** | **1.00** | **Statistically non-significant** |
| | **CellSize_15Å - CellSize_20Å** | **1.00** | **Statistically non-significant** |
| **$R_g$** Total (1001*3) =3003 datapoints | CellSize_3Å - CellSize_5Å | 0.00 | Statistically significant |
| | CellSize_3Å - CellSize_10Å | 0.00 | Statistically significant |
| | CellSize_3Å - CellSize_15Å | 0.00 | Statistically significant |
| | CellSize_3Å - CellSize_20Å | 0.00 | Statistically significant |
| | CellSize_5Å - CellSize_10Å | 0.00 | Statistically significant |
| | CellSize_5Å - CellSize_15Å | 0.00 | Statistically significant |
| | CellSize_5Å - CellSize_20Å | 0.00 | Statistically significant |
| | CellSize_10Å - CellSize_15Å | 0.00 | Statistically significant |
| | CellSize_10Å - CellSize_20Å | 0.00 | Statistically significant |
| | **CellSize_15Å - CellSize_20Å** | **1.00** | **Statistically non-significant** |
| **SASA** Total (1001*3) =3003 datapoints | CellSize_3Å - CellSize_5Å | 0.00 | Statistically significant |
| | CellSize_3Å - CellSize_10Å | 0.00 | Statistically significant |
| | CellSize_3Å - CellSize_15Å | 0.00 | Statistically significant |
| | CellSize_3Å - CellSize_20Å | 0.00 | Statistically significant |
| | CellSize_5Å - CellSize_10Å | 0.00 | Statistically significant |
| | CellSize_5Å - CellSize_15Å | 0.00 | Statistically significant |
| | CellSize_5Å - CellSize_20Å | 0.00 | Statistically significant |
| | CellSize_10Å - CellSize_15Å | 0.00 | Statistically significant |
| | CellSize_10Å - CellSize_20Å | 0.00 | Statistically significant |
| | **CellSize_15Å - CellSize_20Å** | **0.66** | **Statistically non-significant** |
| **PSA** | CellSize_3Å - CellSize_5Å | 0.04 | Statistically significant |
| | CellSize_3Å - CellSize_10Å | 0.00 | Statistically significant |

| | | | |
|---|---|---|---|
| Total (1001*3) =3003 datapoints | CellSize_3Å - CellSize_15Å | 0.00 | Statistically significant |
| | CellSize_3Å - CellSize_20Å | 0.00 | Statistically significant |
| | CellSize_5Å - CellSize_10Å | 0.00 | Statistically significant |
| | CellSize_5Å - CellSize_15Å | 0.00 | Statistically significant |
| | CellSize_5Å - CellSize_20Å | 0.00 | Statistically significant |
| | CellSize_10Å - CellSize_15Å | 0.05 | Statistically significant |
| | **CellSize_10Å - CellSize_20Å** | **0.66** | **Statistically non-significant** |
| | **CellSize_15Å - CellSize_20Å** | **1.00** | **Statistically non-significant** |
| **RMSF per atom** Total (1001*3) =3003 datapoints | CellSize_3Å - CellSize_5Å | 0.00 | Statistically significant |
| | CellSize_3Å - CellSize_10Å | 0.00 | Statistically significant |
| | CellSize_3Å - CellSize_15Å | 0.00 | Statistically significant |
| | CellSize_3Å - CellSize_20Å | 0.00 | Statistically significant |
| | CellSize_5Å - CellSize_10Å | 0.00 | Statistically significant |
| | CellSize_5Å - CellSize_15Å | 0.00 | Statistically significant |
| | CellSize_5Å - CellSize_20Å | 0.00 | Statistically significant |
| | CellSize_10Å - CellSize_15Å | 0.00 | Statistically significant |
| | CellSize_10Å - CellSize_20Å | 0.00 | Statistically significant |
| | **CellSize_15Å - CellSize_20Å** | **1.00** | **Statistically non-significant** |
| **RMSF per residue** Total (1001*3) =3003 datapoints | CellSize_3Å - CellSize_5Å | 0.00 | Statistically significant |
| | CellSize_3Å - CellSize_10Å | 0.00 | Statistically significant |
| | CellSize_3Å - CellSize_15Å | 0.00 | Statistically significant |
| | CellSize_3Å - CellSize_20Å | 0.00 | Statistically significant |
| | CellSize_5Å - CellSize_10Å | 0.05 | Statistically significant |
| | CellSize_5Å - CellSize_15Å | 0.00 | Statistically significant |
| | CellSize_5Å - CellSize_20Å | 0.00 | Statistically significant |
| | **CellSize_10Å - CellSize_15Å** | **1.00** | **Statistically non-significant** |
| | **CellSize_10Å - CellSize_20Å** | **1.00** | **Statistically non-significant** |
| | **CellSize_15Å - CellSize_20Å** | **1.00** | **Statistically non-significant** |
| **Nucleotide-nucleotide hydrogen bonds** Total (500*3) =1500 datapoints | CellSize_3Å - CellSize_5Å | 0.01 | Statistically significant |
| | CellSize_3Å - CellSize_10Å | 0.00 | Statistically significant |
| | CellSize_3Å - CellSize_15Å | 0.00 | Statistically significant |
| | CellSize_3Å - CellSize_20Å | 0.00 | Statistically significant |
| | CellSize_5Å - CellSize_10Å | 0.00 | Statistically significant |
| | CellSize_5Å - CellSize_15Å | 0.00 | Statistically significant |
| | CellSize_5Å - CellSize_20Å | 0.00 | Statistically significant |
| | CellSize_10Å - CellSize_15Å | 0.00 | Statistically significant |

| | | | |
|---|---|---|---|
| | CellSize_10Å - CellSize_20Å | 0.00 | Statistically significant |
| | CellSize_15Å - CellSize_20Å | 0.03 | Statistically significant |
| **Nucleotide-nucleotide π-π stacking interaction** Total (500*3) =1500 datapoints | CellSize_3Å - CellSize_5Å | 0.00 | Statistically significant |
| | CellSize_3Å - CellSize_10Å | 0.00 | Statistically significant |
| | CellSize_3Å - CellSize_15Å | 0.00 | Statistically significant |
| | CellSize_3Å - CellSize_20Å | 0.00 | Statistically significant |
| | CellSize_5Å - CellSize_10Å | 0.08 | Statistically insignificant |
| | CellSize_5Å - CellSize_15Å | 0.01 | Statistically significant |
| | CellSize_5Å - CellSize_20Å | 0.00 | Statistically significant |
| | **CellSize_10Å - CellSize_15Å** | **1.00** | **Statistically non-significant** |
| | CellSize_10Å - CellSize_20Å | 0.00 | Statistically significant |
| | CellSize_15Å - CellSize_20Å | 0.01 | Statistically significant |
| **Nucleotide-water hydrogen bonds** Total (500*3) =1500 datapoints | CellSize_3Å - CellSize_5Å | 0.00 | Statistically significant |
| | CellSize_3Å - CellSize_10Å | 0.00 | Statistically significant |
| | CellSize_3Å - CellSize_15Å | 0.00 | Statistically significant |
| | CellSize_3Å - CellSize_20Å | 0.00 | Statistically significant |
| | CellSize_5Å - CellSize_10Å | 0.00 | Statistically significant |
| | CellSize_5Å - CellSize_15Å | 0.00 | Statistically significant |
| | CellSize_5Å - CellSize_20Å | 0.00 | Statistically significant |
| | **CellSize_10Å - CellSize_15Å** | **1.00** | **Statistically non-significant** |
| | **CellSize_10Å - CellSize_20Å** | **1.00** | **Statistically non-significant** |
| | **CellSize_15Å - CellSize_20Å** | **1.00** | **Statistically non-significant** |
| **Monomer 1 (Antisense strand)** | | | |
| **RMSD** Total (1001*3) =3003 datapoints | CellSize_3Å - CellSize_5Å | 0.00 | Statistically significant |
| | CellSize_3Å - CellSize_10Å | 0.00 | Statistically significant |
| | CellSize_3Å - CellSize_15Å | 0.00 | Statistically significant |
| | CellSize_3Å - CellSize_20Å | 0.00 | Statistically significant |
| | **CellSize_5Å - CellSize_10Å** | **1.00** | **Statistically non-significant** |
| | CellSize_5Å - CellSize_15Å | 0.00 | Statistically significant |
| | CellSize_5Å - CellSize_20Å | 0.00 | Statistically significant |
| | CellSize_10Å - CellSize_15Å | 0.00 | Statistically significant |
| | CellSize_10Å - CellSize_20Å | 0.00 | Statistically significant |
| | **CellSize_15Å - CellSize_20Å** | 0.00 | Statistically significant |
| $\mathbf{R_g}$ | CellSize_3Å - CellSize_5Å | 0.00 | Statistically significant |
| | CellSize_3Å - CellSize_10Å | 0.00 | Statistically significant |
| | CellSize_3Å - CellSize_15Å | 0.00 | Statistically significant |

| | | | |
|---|---|---|---|
| Total (1001*3) =3003 datapoints | CellSize_3Å - CellSize_20Å | 0.00 | Statistically significant |
| | CellSize_5Å - CellSize_10Å | 0.00 | Statistically significant |
| | CellSize_5Å - CellSize_15Å | 0.00 | Statistically significant |
| | CellSize_5Å - CellSize_20Å | 0.00 | Statistically significant |
| | **CellSize_10Å - CellSize_15Å** | **1.00** | **Statistically non-significant** |
| | CellSize_10Å - CellSize_20Å | 0.00 | Statistically significant |
| | CellSize_15Å - CellSize_20Å | 0.00 | Statistically significant |
| **SASA** Total (1001*3) =3003 datapoints | CellSize_3Å - CellSize_5Å | 0.00 | Statistically significant |
| | CellSize_3Å - CellSize_10Å | 0.00 | Statistically significant |
| | CellSize_3Å - CellSize_15Å | 0.00 | Statistically significant |
| | CellSize_3Å - CellSize_20Å | 0.00 | Statistically significant |
| | CellSize_5Å - CellSize_10Å | 0.00 | Statistically significant |
| | CellSize_5Å - CellSize_15Å | 0.00 | Statistically significant |
| | CellSize_5Å - CellSize_20Å | 0.00 | Statistically significant |
| | CellSize_10Å - CellSize_15Å | 0.00 | Statistically significant |
| | CellSize_10Å - CellSize_20Å | 0.00 | Statistically significant |
| | CellSize_15Å - CellSize_20Å | 0.00 | Statistically significant |
| **PSA** Total (1001*3) =3003 datapoints | CellSize_3Å - CellSize_5Å | 0.00 | Statistically significant |
| | CellSize_3Å - CellSize_10Å | 0.00 | Statistically significant |
| | CellSize_3Å - CellSize_15Å | 0.00 | Statistically significant |
| | CellSize_3Å - CellSize_20Å | 0.00 | Statistically significant |
| | CellSize_5Å - CellSize_10Å | 0.00 | Statistically significant |
| | CellSize_5Å - CellSize_15Å | 0.00 | Statistically significant |
| | CellSize_5Å - CellSize_20Å | 0.00 | Statistically significant |
| | CellSize_10Å - CellSize_15Å | 0.00 | Statistically significant |
| | CellSize_10Å - CellSize_20Å | 0.00 | Statistically significant |
| | CellSize_15Å - CellSize_20Å | 0.00 | Statistically significant |
| **RMSF per atom** Total (1001*3) =3003 datapoints | CellSize_3Å - CellSize_5Å | 0.00 | Statistically significant |
| | **CellSize_3Å - CellSize_10Å** | **0.08** | **Statistically non-significant** |
| | **CellSize_3Å - CellSize_15Å** | **1.00** | **Statistically non-significant** |
| | CellSize_3Å - CellSize_20Å | 0.00 | Statistically significant |
| | CellSize_5Å - CellSize_10Å | 0.00 | Statistically significant |
| | CellSize_5Å - CellSize_15Å | 0.00 | Statistically significant |
| | CellSize_5Å - CellSize_20Å | 0.00 | Statistically significant |
| | CellSize_10Å - CellSize_15Å | 0.00 | Statistically significant |
| | CellSize_10Å - CellSize_20Å | 0.00 | Statistically significant |

| | | | |
|---|---|---|---|
| | CellSize_15Å - CellSize_20Å | 0.00 | Statistically significant |
| **RMSF per residue** Total (1001*3) =3003 datapoints | **CellSize_3Å - CellSize_5Å** | **0.17** | **Statistically non-significant** |
| | **CellSize_3Å - CellSize_10Å** | **1.00** | **Statistically non-significant** |
| | **CellSize_3Å - CellSize_15Å** | **1.00** | **Statistically non-significant** |
| | **CellSize_3Å - CellSize_20Å** | **1.00** | **Statistically non-significant** |
| | **CellSize_5Å - CellSize_10Å** | **0.16** | **Statistically non-significant** |
| | CellSize_5Å - CellSize_15Å | 0.01 | Statistically significant |
| | CellSize_5Å - CellSize_20Å | 0.00 | Statistically significant |
| | **CellSize_10Å - CellSize_15Å** | **1.00** | **Statistically non-significant** |
| | **CellSize_10Å - CellSize_20Å** | **1.00** | **Statistically non-significant** |
| | **CellSize_15Å - CellSize_20Å** | **1.00** | **Statistically non-significant** |
| **Nucleotide-nucleotide hydrogen bonds** Total (500*3) =1500 datapoints | CellSize_3Å - CellSize_5Å | 0.00 | Statistically significant |
| | CellSize_3Å - CellSize_10Å | 0.00 | Statistically significant |
| | CellSize_3Å - CellSize_15Å | 0.00 | Statistically significant |
| | CellSize_3Å - CellSize_20Å | 0.00 | Statistically significant |
| | CellSize_5Å - CellSize_10Å | 0.00 | Statistically significant |
| | CellSize_5Å - CellSize_15Å | 0.00 | Statistically significant |
| | **CellSize_5Å - CellSize_20Å** | **1.00** | **Statistically non-significant** |
| | CellSize_10Å - CellSize_15Å | 0.00 | Statistically significant |
| | CellSize_10Å - CellSize_20Å | 0.00 | Statistically significant |
| | CellSize_15Å - CellSize_20Å | 0.00 | Statistically significant |
| **Nucleotide-nucleotide *π-π* stacking interaction** Total (500*3) =1500 datapoints | CellSize_3Å - CellSize_5Å | 0.00 | Statistically significant |
| | CellSize_3Å - CellSize_10Å | 0.00 | Statistically significant |
| | CellSize_3Å - CellSize_15Å | 0.00 | Statistically significant |
| | CellSize_3Å - CellSize_20Å | 0.00 | Statistically significant |
| | CellSize_5Å - CellSize_10Å | 0.00 | Statistically significant |
| | CellSize_5Å - CellSize_15Å | 0.00 | Statistically significant |
| | **CellSize_5Å - CellSize_20Å** | **1.00** | **Statistically non-significant** |
| | CellSize_10Å - CellSize_15Å | 0.00 | Statistically significant |
| | CellSize_10Å - CellSize_20Å | 0.00 | Statistically significant |
| | CellSize_15Å - CellSize_20Å | 0.00 | Statistically significant |
| **Nucleotide-water hydrogen bonds** | CellSize_3Å - CellSize_5Å | 0.00 | Statistically significant |
| | CellSize_3Å - CellSize_10Å | 0.00 | Statistically significant |
| | CellSize_3Å - CellSize_15Å | 0.00 | Statistically significant |
| | **CellSize_3Å - CellSize_20Å** | **1.00** | **Statistically non-significant** |
| | CellSize_5Å - CellSize_10Å | 0.00 | Statistically significant |

| | | | |
|---|---|---|---|
| Total (500*3) =1500 datapoints | CellSize_5Å - CellSize_15Å | 0.00 | Statistically significant |
| | CellSize_5Å - CellSize_20Å | 0.00 | Statistically significant |
| | **CellSize_10Å - CellSize_15Å** | **1.00** | **Statistically non-significant** |
| | CellSize_10Å - CellSize_20Å | 0.00 | Statistically significant |
| | CellSize_15Å - CellSize_20Å | 0.00 | Statistically significant |
| **Monomer 2 (Sense strand)** | | | |
| **RMSD** Total (1001*3) =3003 datapoints | **CellSize_3Å - CellSize_5Å** | **1.00** | **Statistically non-significant** |
| | CellSize_3Å - CellSize_10Å | 0.00 | Statistically significant |
| | CellSize_3Å - CellSize_15Å | 0.05 | Statistically significant |
| | CellSize_3Å - CellSize_20Å | 0.00 | Statistically significant |
| | CellSize_5Å - CellSize_10Å | 0.00 | Statistically significant |
| | CellSize_5Å - CellSize_15Å | 0.00 | Statistically significant |
| | CellSize_5Å - CellSize_20Å | 0.00 | Statistically significant |
| | CellSize_10Å - CellSize_15Å | 0.00 | Statistically significant |
| | CellSize_10Å - CellSize_20Å | 0.00 | Statistically significant |
| | **CellSize_15Å - CellSize_20Å** | 0.00 | Statistically significant |
| **$R_g$** Total (1001*3) =3003 datapoints | **CellSize_3Å - CellSize_5Å** | **0.05** | **Statistically non-significant** |
| | CellSize_3Å - CellSize_10Å | 0.00 | Statistically significant |
| | **CellSize_3Å - CellSize_15Å** | **1.00** | **Statistically non-significant** |
| | CellSize_3Å - CellSize_20Å | 0.00 | Statistically significant |
| | CellSize_5Å - CellSize_10Å | 0.00 | Statistically significant |
| | **CellSize_5Å - CellSize_15Å** | **0.49** | **Statistically non-significant** |
| | CellSize_5Å - CellSize_20Å | 0.00 | Statistically significant |
| | CellSize_10Å - CellSize_15Å | 0.00 | Statistically significant |
| | CellSize_10Å - CellSize_20Å | 0.00 | Statistically significant |
| | CellSize_15Å - CellSize_20Å | 0.00 | Statistically significant |
| **SASA** Total (1001*3) =3003 datapoints | CellSize_3Å - CellSize_5Å | 0.00 | Statistically significant |
| | CellSize_3Å - CellSize_10Å | 0.00 | Statistically significant |
| | CellSize_3Å - CellSize_15Å | 0.00 | Statistically significant |
| | CellSize_3Å - CellSize_20Å | 0.00 | Statistically significant |
| | CellSize_5Å - CellSize_10Å | 0.00 | Statistically significant |
| | CellSize_5Å - CellSize_15Å | 0.00 | Statistically significant |
| | CellSize_5Å - CellSize_20Å | 0.00 | Statistically significant |
| | CellSize_10Å - CellSize_15Å | 0.00 | Statistically significant |
| | CellSize_10Å - CellSize_20Å | 0.00 | Statistically significant |
| | **CellSize_15Å - CellSize_20Å** | **0.12** | **Statistically non-significant** |

| | | | |
|---|---|---|---|
| **PSA** Total (1001*3) =3003 datapoints | CellSize_3Å - CellSize_5Å | 0.00 | Statistically significant |
| | CellSize_3Å - CellSize_10Å | 0.00 | Statistically significant |
| | CellSize_3Å - CellSize_15Å | 0.00 | Statistically significant |
| | CellSize_3Å - CellSize_20Å | 0.00 | Statistically significant |
| | CellSize_5Å - CellSize_10Å | 0.00 | Statistically significant |
| | CellSize_5Å - CellSize_15Å | 0.00 | Statistically significant |
| | CellSize_5Å - CellSize_20Å | 0.00 | Statistically significant |
| | CellSize_10Å - CellSize_15Å | 0.00 | Statistically significant |
| | CellSize_10Å - CellSize_20Å | 0.00 | Statistically significant |
| | **CellSize_15Å - CellSize_20Å** | **1.00** | **Statistically non-significant** |
| **RMSF per atom** Total (1001*3) =3003 datapoints | CellSize_3Å - CellSize_5Å | 0.00 | Statistically significant |
| | CellSize_3Å - CellSize_10Å | 0.00 | Statistically significant |
| | CellSize_3Å - CellSize_15Å | 0.00 | Statistically significant |
| | **CellSize_3Å - CellSize_20Å** | **0.82** | **Statistically non-significant** |
| | CellSize_5Å - CellSize_10Å | 0.00 | Statistically significant |
| | CellSize_5Å - CellSize_15Å | 0.00 | Statistically significant |
| | CellSize_5Å - CellSize_20Å | 0.00 | Statistically significant |
| | CellSize_10Å - CellSize_15Å | 0.00 | Statistically significant |
| | CellSize_10Å - CellSize_20Å | 0.00 | Statistically significant |
| | **CellSize_15Å - CellSize_20Å** | **0.59** | **Statistically non-significant** |
| **RMSF per residue** Total (1001*3) =3003 datapoints | **CellSize_3Å - CellSize_5Å** | **1.00** | **Statistically non-significant** |
| | **CellSize_3Å - CellSize_10Å** | **0.08** | **Statistically non-significant** |
| | **CellSize_3Å - CellSize_15Å** | **1.00** | **Statistically non-significant** |
| | **CellSize_3Å - CellSize_20Å** | **1.00** | **Statistically non-significant** |
| | CellSize_5Å - CellSize_10Å | 0.01 | Statistically significant |
| | **CellSize_5Å - CellSize_15Å** | **1.00** | **Statistically non-significant** |
| | **CellSize_5Å - CellSize_20Å** | **1.00** | **Statistically non-significant** |
| | CellSize_10Å - CellSize_15Å | 0.00 | Statistically significant |
| | **CellSize_10Å - CellSize_20Å** | **0.29** | **Statistically non-significant** |
| | **CellSize_15Å - CellSize_20Å** | **1.00** | **Statistically non-significant** |
| **Nucleotide-nucleotide hydrogen bonds** Total (500*3) =1500datapoints | CellSize_3Å - CellSize_5Å | 0.00 | Statistically significant |
| | CellSize_3Å - CellSize_10Å | 0.02 | Statistically significant |
| | CellSize_3Å - CellSize_15Å | 0.00 | Statistically significant |
| | CellSize_3Å - CellSize_20Å | 0.00 | Statistically significant |
| | CellSize_5Å - CellSize_10Å | 0.00 | Statistically significant |
| | CellSize_5Å - CellSize_15Å | 0.00 | Statistically significant |

| | | | |
|---|---|---|---|
| | CellSize_5Å - CellSize_20Å | 0.00 | Statistically significant |
| | **CellSize_10Å - CellSize_15Å** | **0.53** | **Statistically non-significant** |
| | CellSize_10Å - CellSize_20Å | 0.00 | Statistically significant |
| | CellSize_15Å - CellSize_20Å | 0.00 | Statistically significant |
| **Nucleotide-nucleotide π-π stacking interaction** Total (500*3) =1500 datapoints | CellSize_3Å - CellSize_5Å | 0.00 | Statistically significant |
| | CellSize_3Å - CellSize_10Å | 0.00 | Statistically significant |
| | CellSize_3Å - CellSize_15Å | 0.00 | Statistically significant |
| | CellSize_3Å - CellSize_20Å | 0.00 | Statistically significant |
| | CellSize_5Å - CellSize_10Å | 0.00 | Statistically significant |
| | CellSize_5Å - CellSize_15Å | 0.00 | Statistically significant |
| | CellSize_5Å - CellSize_20Å | 0.00 | Statistically significant |
| | CellSize_10Å - CellSize_15Å | 0.00 | Statistically significant |
| | **CellSize_10Å - CellSize_20Å** | **0.26** | **Statistically non-significant** |
| | CellSize_15Å - CellSize_20Å | 0.00 | Statistically significant |
| **Nucleotide-water hydrogen bonds** Total (500*3) =1500 datapoints | CellSize_3Å - CellSize_5Å | 0.00 | Statistically significant |
| | CellSize_3Å - CellSize_10Å | 0.00 | Statistically significant |
| | CellSize_3Å - CellSize_15Å | 0.00 | Statistically significant |
| | CellSize_3Å - CellSize_20Å | 0.00 | Statistically significant |
| | CellSize_5Å - CellSize_10Å | 0.00 | Statistically significant |
| | CellSize_5Å - CellSize_15Å | 0.01 | Statistically significant |
| | CellSize_5Å - CellSize_20Å | 0.00 | Statistically significant |
| | CellSize_10Å - CellSize_15Å | 0.00 | Statistically significant |
| | **CellSize_10Å - CellSize_20Å** | **1.00** | **Statistically non-significant** |
| | CellSize_15Å - CellSize_20Å | 0.00 | Statistically significant |

**Table S7. The distance (Å) between the 5'- and 3'-ends of DNA is measured from each representative structure of the MD replicate.**

| | System | Strand and residue | | | Distance |
|---|---|---|---|---|---|
| **PDB: 7EDS** | | Antisense strand | 5' Adenine | 3' Guanine | 101.90 Å |
| | | Sense strand | 3' Thymine | 5' Cytosine | 102.36 Å |
| **Dimer (Double-stranded)** | | | | | |
| **3 Å** | **MD1** | Antisense strand | 5' Adenine | 3' Guanine | 107.19 Å |
| | | Sense strand | 3' Thymine | 5' Cytosine | 106.13 Å |
| | **MD2** | Antisense strand | 5' Adenine | 3' Guanine | 108.80 Å |
| | | Sense strand | 3' Thymine | 5' Cytosine | 103.66Å |
| | **MD3** | Antisense strand | 5' Adenine | 3' Guanine | 102.50Å |
| | | Sense strand | 3' Thymine | 5' Cytosine | 100.35 Å |
| **5 Å** | **MD1** | Antisense strand | 5' Adenine | 3' Guanine | 98.15 Å |
| | | Sense strand | 3' Thymine | 5' Cytosine | 108.65 Å |
| | **MD2** | Antisense strand | 5' Adenine | 3' Guanine | 103.37 Å |
| | | Sense strand | 3' Thymine | 5' Cytosine | 106.39 Å |
| | **MD3** | Antisense strand | 5' Adenine | 3' Guanine | 106.53 Å |
| | | Sense strand | 3' Thymine | 5' Cytosine | 99.68 Å |
| **10 Å** | **MD1** | Antisense strand | 5' Adenine | 3' Guanine | 104.02 Å |
| | | Sense strand | 3' Thymine | 5' Cytosine | 99.84 Å |
| | **MD2** | Antisense strand | 5' Adenine | 3' Guanine | 106.91 Å |
| | | Sense strand | 3' Thymine | 5' Cytosine | 97.87 Å |
| | **MD3** | Antisense strand | 5' Adenine | 3' Guanine | 105.30 Å |
| | | Sense strand | 3' Thymine | 5' Cytosine | 95.99 Å |
| **15 Å** | **MD1** | Antisense strand | 5' Adenine | 3' Guanine | 103.72 Å |
| | | Sense strand | 3' Thymine | 5' Cytosine | 105.92 Å |
| | **MD2** | Antisense strand | 5' Adenine | 3' Guanine | 106.77 Å |
| | | Sense strand | 3' Thymine | 5' Cytosine | 99.42 Å |
| | **MD3** | Antisense strand | 5' Adenine | 3' Guanine | 100.01 Å |
| | | Sense strand | 3' Thymine | 5' Cytosine | 96.68 Å |
| **20 Å** | **MD1** | Antisense strand | 5' Adenine | 3' Guanine | 94.18 Å |
| | | Sense strand | 3' Thymine | 5' Cytosine | 103.38 Å |
| | **MD2** | Antisense strand | 5' Adenine | 3' Guanine | 101.62 Å |
| | | Sense strand | 3' Thymine | 5' Cytosine | 101.55 Å |
| | **MD3** | Antisense strand | 5' Adenine | 3' Guanine | 107.09 Å |
| | | Sense strand | 3' Thymine | 5' Cytosine | 99.05 Å |
| **Monomer 1 (Antisense strand)** | | | | | |
| 3 Å | **MD1** | Antisense strand | 5' Adenine | 3' Guanine | 54.77 Å |
| | **MD2** | | | | 53.55 Å |
| | **MD3** | | | | 89.80 Å |
| 5 Å | **MD1** | | | | 66.87 Å |
| | **MD2** | | | | 28.39 Å |
| | **MD3** | | | | 31.95 Å |

| | | | | | |
|---|---|---|---|---|---|
| 10 Å | **MD1** | | | | 33.34 Å |
| | **MD2** | | | | 60.65 Å |
| | **MD3** | | | | 28.50 Å |
| **15 Å** | **MD1** | | | | 75.21 Å |
| | **MD2** | | | | 99.50 Å |
| | **MD3** | | | | 25.17 Å |
| **20 Å** | **MD1** | | | | 16.14 Å |
| | **MD2** | | | | 34.69 Å |
| | **MD3** | | | | 36.62 Å |
| **Monomer 2 (Sense strand)** | | | | | |
| **3 Å** | **MD1** | Sense strand | 3' Thymine | 5' Cytosine | 84.68 Å |
| | **MD2** | | | | 47.20 Å |
| | **MD3** | | | | 59.11 Å |
| **5 Å** | **MD1** | | | | 92.79 Å |
| | **MD2** | | | | 63.14 Å |
| | **MD3** | | | | 39.10 Å |
| **10 Å** | **MD1** | | | | 40.24 Å |
| | **MD2** | | | | 62.62 Å |
| | **MD3** | | | | 49.16 Å |
| **15 Å** | **MD1** | | | | 37.72 Å |
| | **MD2** | | | | 40.97 Å |
| | **MD3** | | | | 54.19 Å |
| **20 Å** | **MD1** | | | | 18.54 Å |
| | **MD2** | | | | 75.23 Å |
| | **MD3** | | | | 45.18 Å |

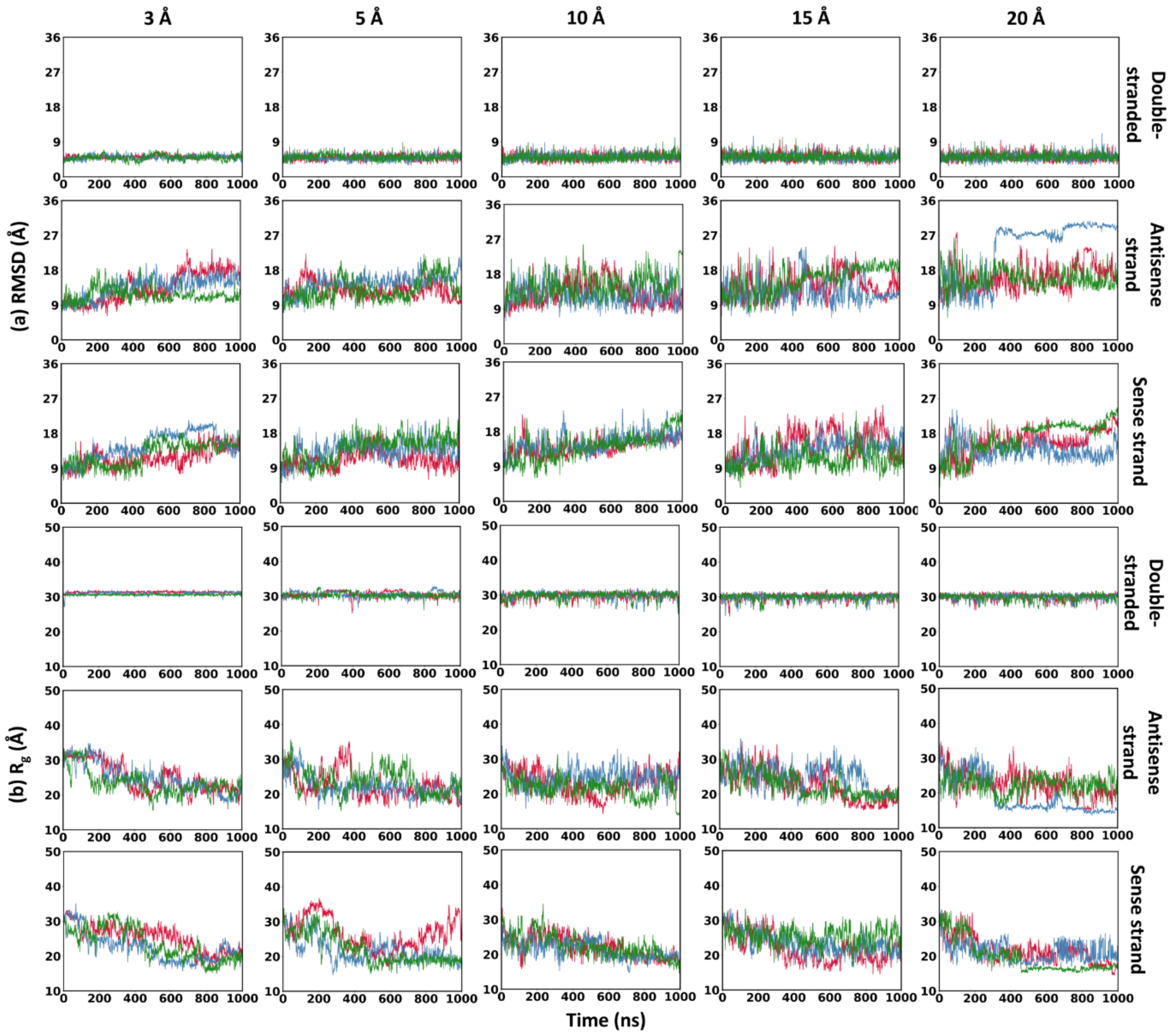


**Figure S1. RMSD and $R_g$ analysis over 1000 ns for the three DNA forms (double-stranded, antisense strand, sense strand) across five periodic simulation cell sizes (3, 5, 10, 15, and 20 Å). Red, blue, and green color represent RUN1, RUN2, and RUN3. (a)** Root Mean Square Deviation (RMSD, Å) relative to the initial structure. **(b)** Radius of gyration ($R_g$, Å).

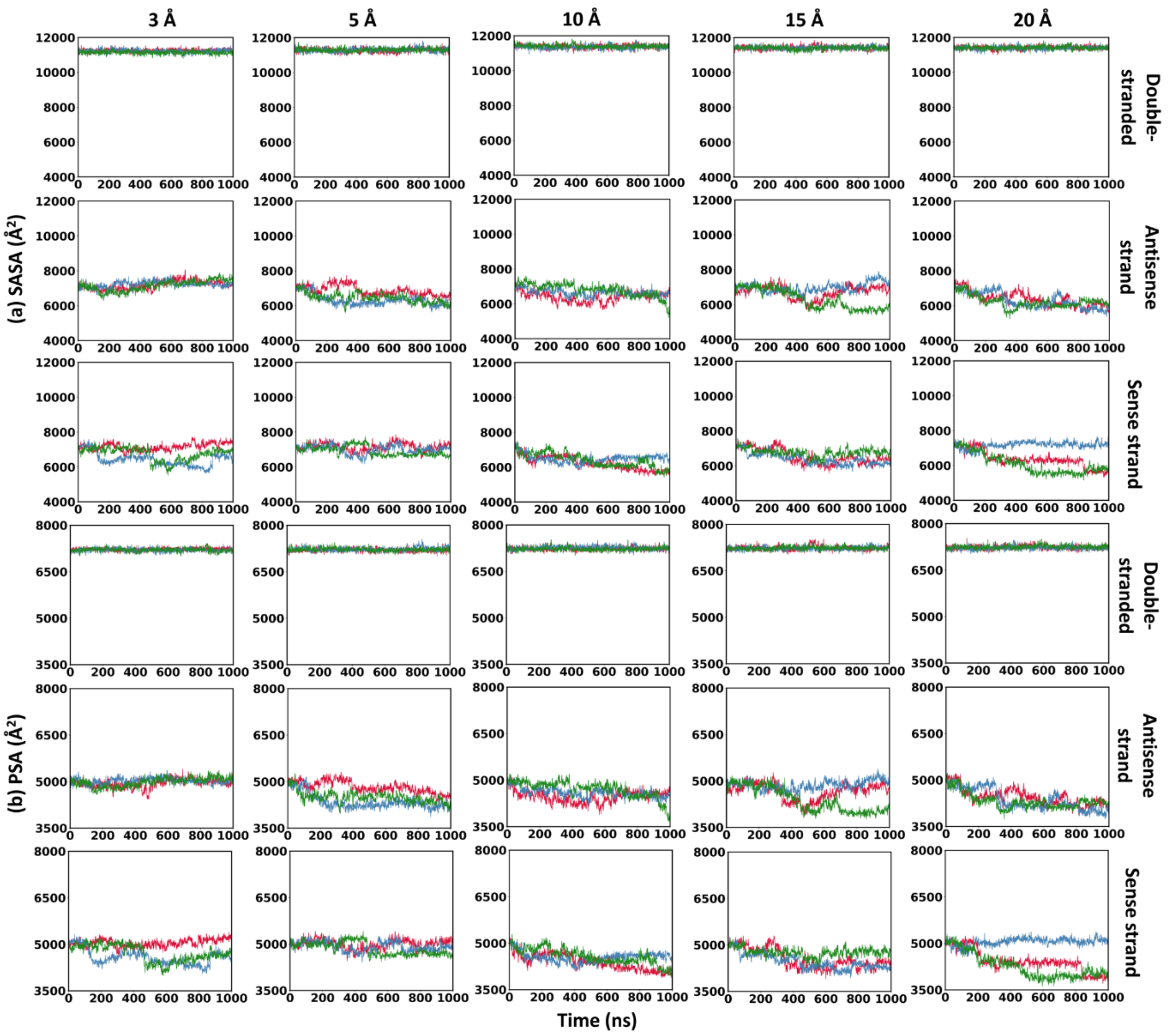


**Figure S2. SASA and PSA analysis over 1000 ns for the three DNA forms (double-stranded, antisense strand, sense strand) across five periodic simulation cell sizes (3, 5, 10, 15, and 20 Å). Red, blue, and green color represent RUN1, RUN2, and RUN3. (a)** Solvent-Accessible Surface Area (SASA, Å²), and **(b)** polar Surface Area (PSA, Å²).

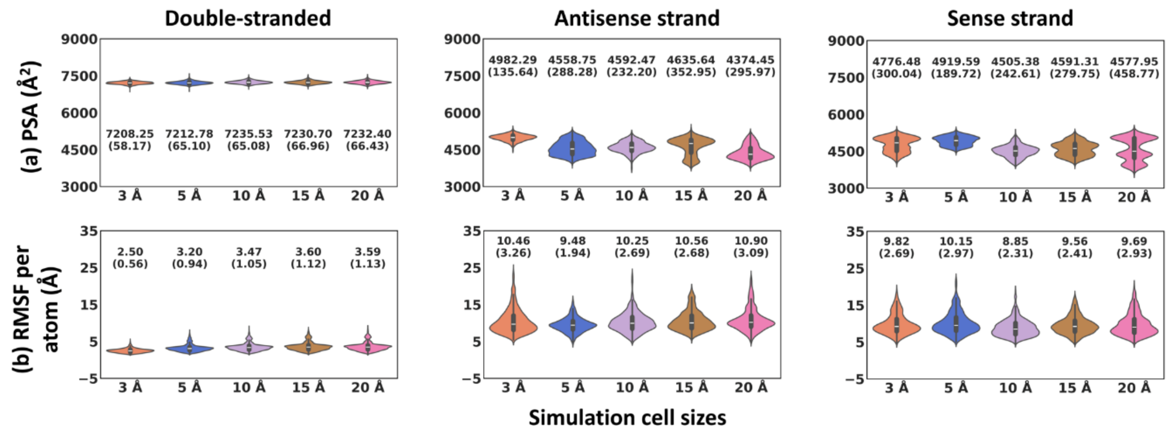


**Figure S3. Violin plots showing the full trajectory distribution of the three DNA forms (double-stranded, antisense strand, Sense strand) across five simulation cell sizes (3, 5, 10, 15, and 20 Å). Each violin represents 3,003 data points compiled from three independent MD replicates. Mean (top annotation) and standard deviation (in brackets) are shown. (a)** Polar Surface Area (PSA, Å²). **(b)** Root Mean Square Fluctuation (RMSF, Å) per atom.

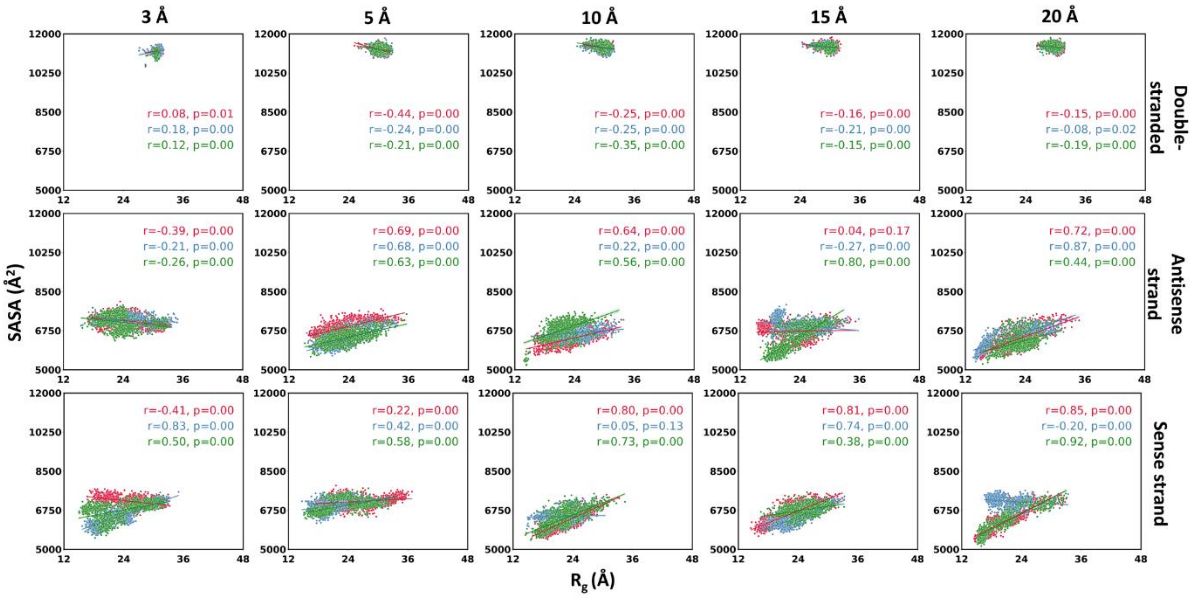


**Figure S4. Pearson correlation plots between radius of gyration ($R_g$) and solvent-accessible surface area (SASA) for the three forms of DNA (double-stranded, antisense strand, sense strand) across five simulation cell sizes (3, 5, 10, 15, and 20 Å). Red, blue, and green color represent RUN1, RUN2, and RUN3.**

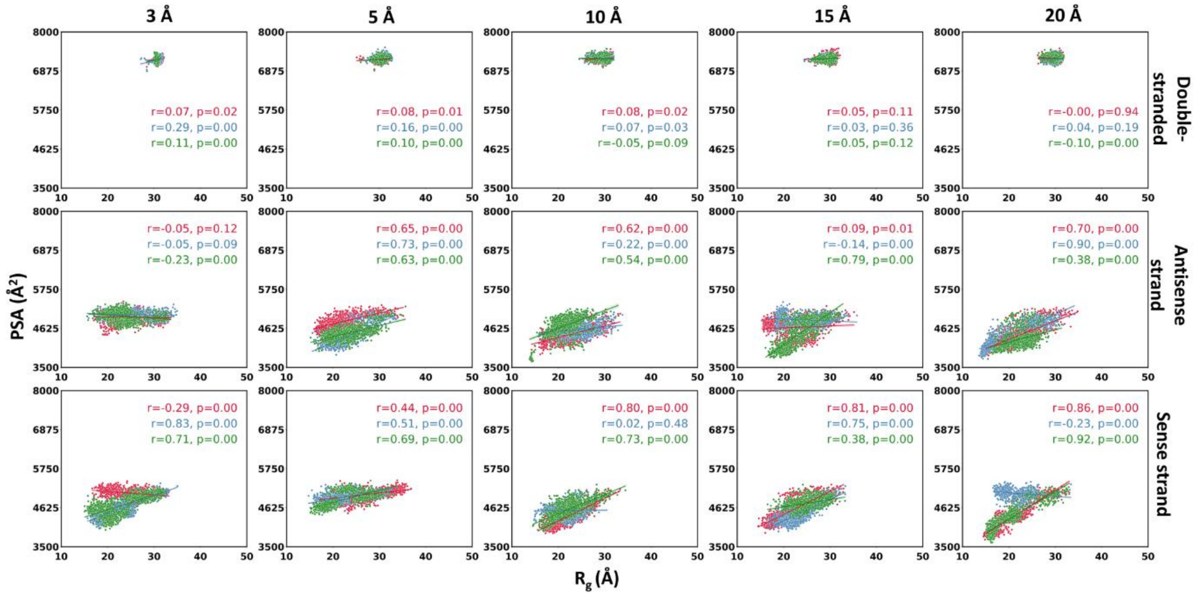


**Figure S5. Pearson correlation plots between radius of gyration ($R_g$) and polar surface area (PSA) for the three forms of DNA (double-stranded, antisense strand, sense strand) across five simulation cell sizes (3, 5, 10, 15, and 20 Å). Red, blue, and green color represent RUN1, RUN2, and RUN3.**

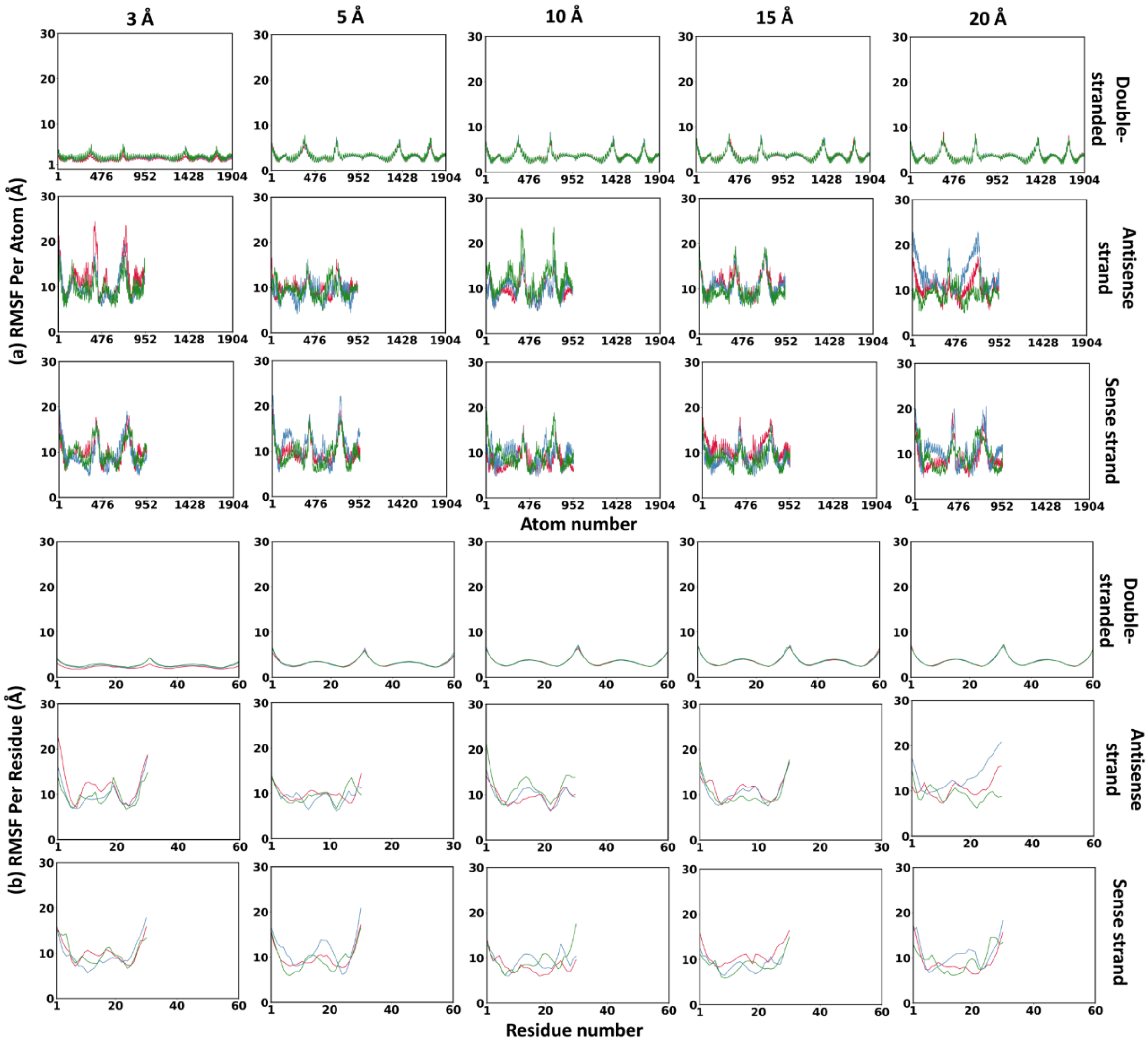


**Figure S6. Per-atom and per-residue fluctuations over 1000 ns for the three DNA forms (double-stranded, antisense strand, sense strand) across five simulation cell sizes (3, 5, 10, 15, and 20 Å). Red, blue, and green color represent RUN1, RUN2, and RUN3. (a)** RMSF per atom (Å), and **(b)** RMSF per residue (Å).

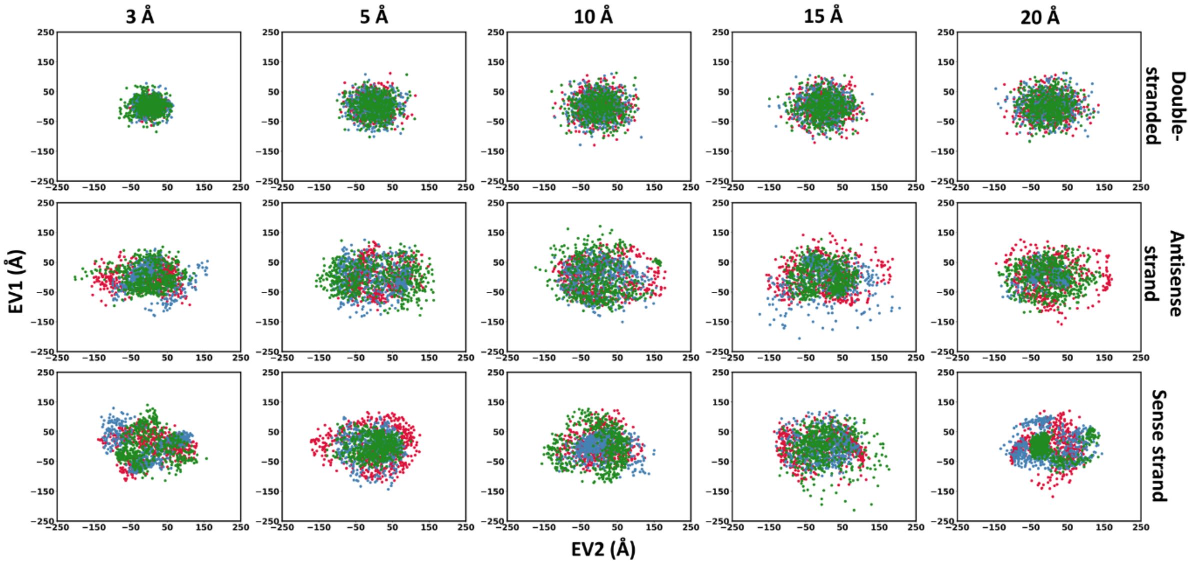


**Figure S7. Principal Component Analysis (PCA) of three DNA forms across five simulation cell sizes (3, 5, 10, 15, and 20 Å). Scatter plots of eigenvector 1 (PC1) versus eigenvector 2 (PC2). Red, blue, and green color represent RUN1, RUN2, and RUN3.**

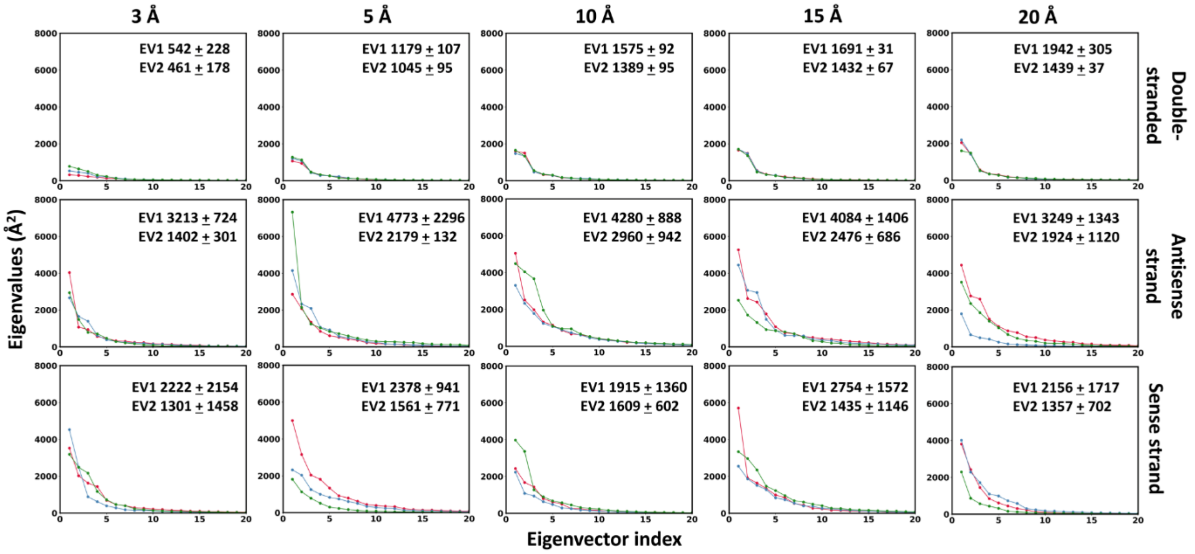


**Figure S8. Principal Component Analysis (PCA) of three DNA forms across five simulation cell sizes (3, 5, 10, 15, and 20 Å). Eigenvalues are plotted against eigenvectors. Red, blue, and green color represent RUN1, RUN2, and RUN3.**

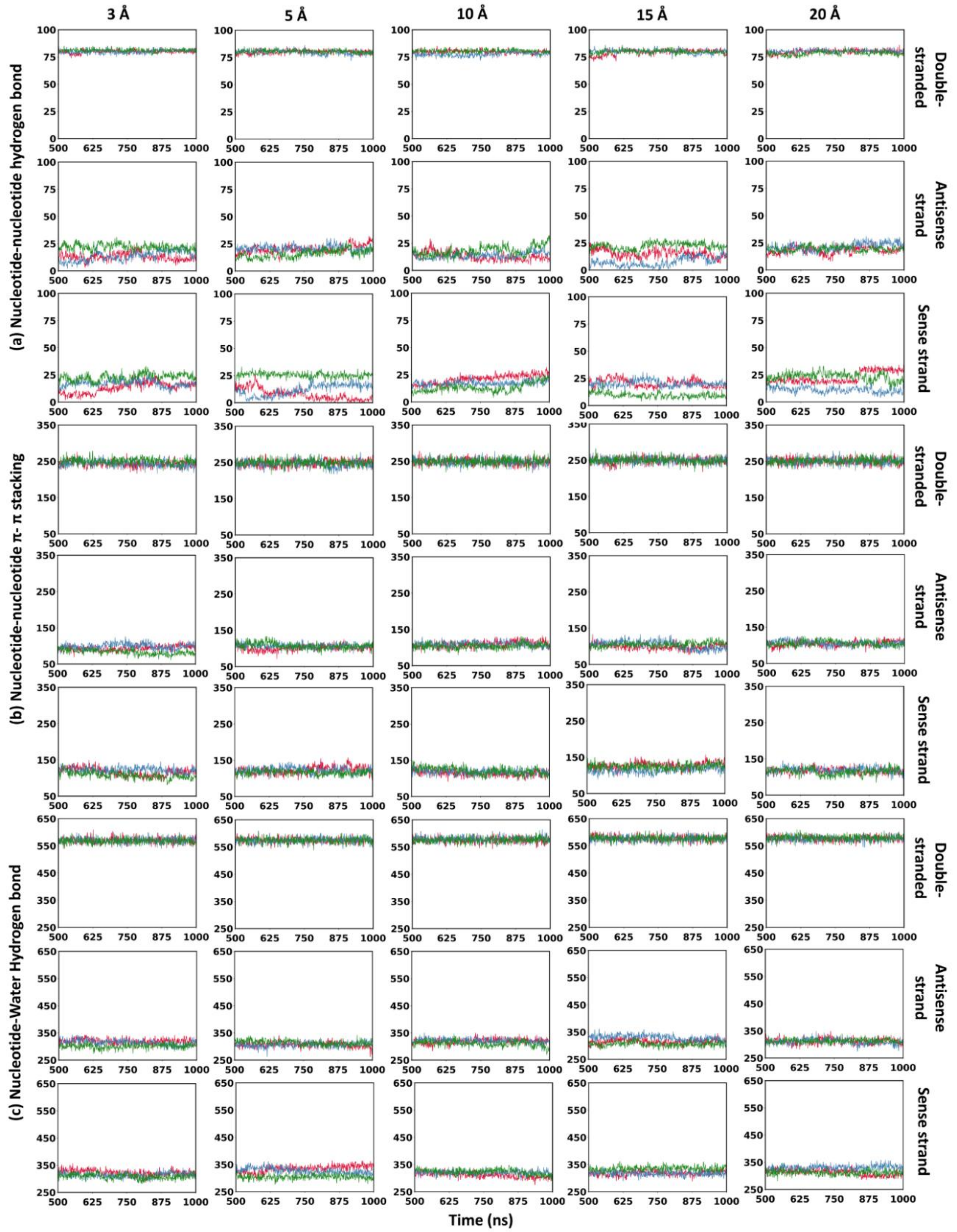


**Figure S9. Analysis of equilibrated trajectory (last 500 ns) for the three DNA forms (double-stranded, antisense strand, sense strand) across five simulation cell sizes (3, 5, 10, 15, and 20 Å). Red, blue, and green color represent RUN1, RUN2, and RUN3. (a)** Nucleotide-nucleotide hydrogen bond count, **(b)** nucleotide-nucleotide π-π stacking, and **(c)** nucleotide-water hydrogen bond.

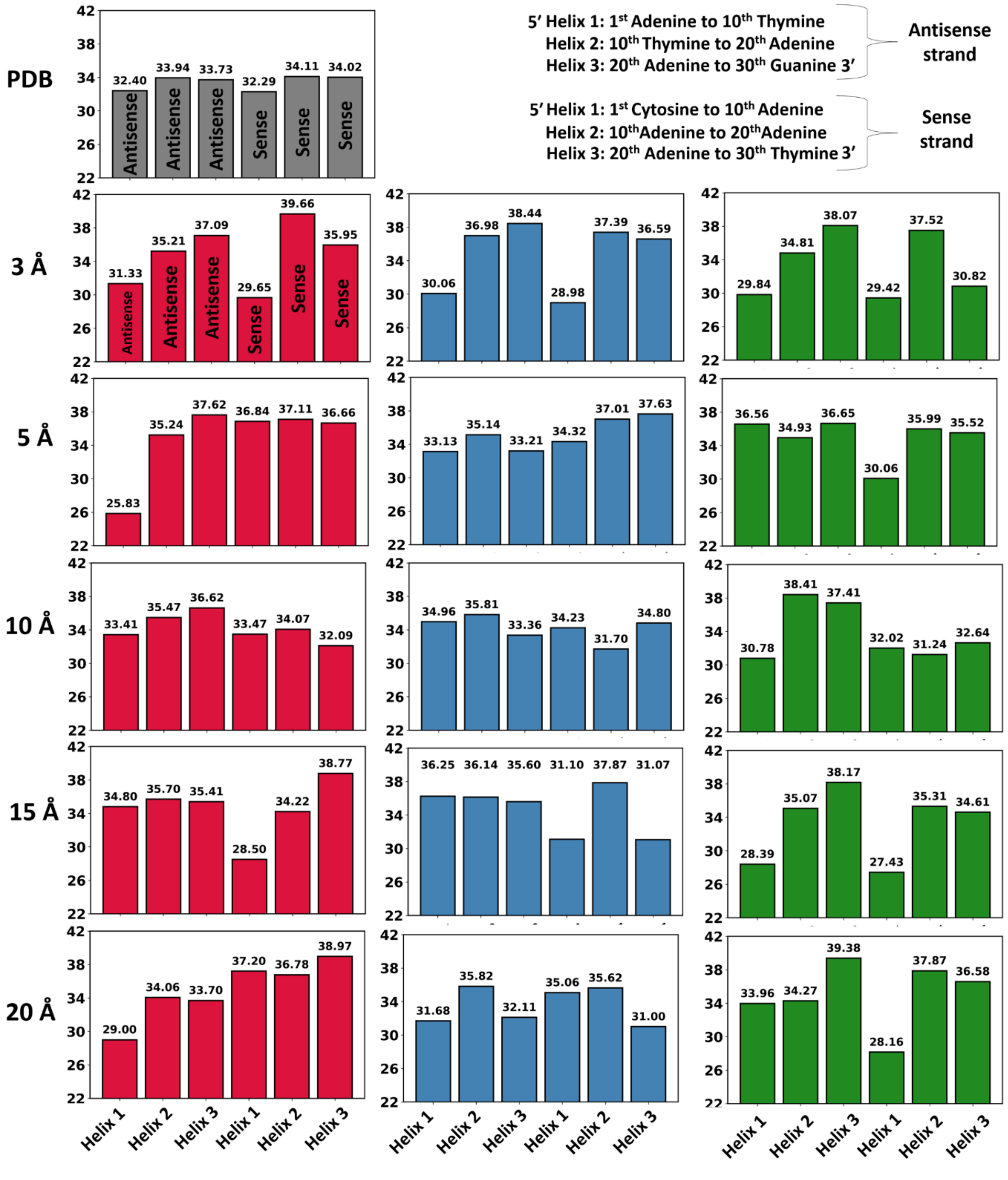


**Figure S10. Helical pitch calculated for double-stranded representative structure across five cell sizes.** Helical pitch distance is mentioned on Y axis and helices are mentioned on X axis. Each strand has three helices sequentially named as helix 1, helix 2 and helix 3 in 5' to 3' direction. Helix 1 (**1st Adenine to 10th Thymine**), helix 2 (**10th Thymine to 20th Adenine**), helix 3 (**20th Adenine to 30 Guanine**) for antisense strand, while helix 1 (**1st Cytosine to 10th Adenine**), helix

2 (**10th Adenine to 20th Adenine**) and helix 3 (**20th Adenine to 30th Thymine**) for sense strand. Grey, red, blue, and green color represent PDB, RUN1, RUN2, and RUN3.

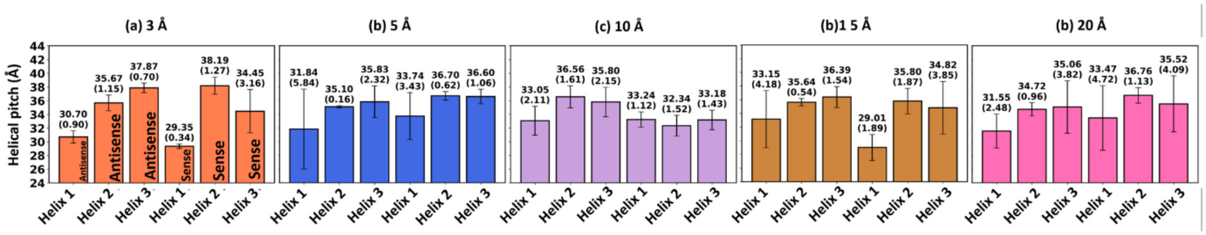


**Figure S11. Average and standard deviation of helical pitch for three runs. Helix pitch is calculated only for the double-stranded representative structures. An annotation represents average (top) and stranded deviation (bracket).**

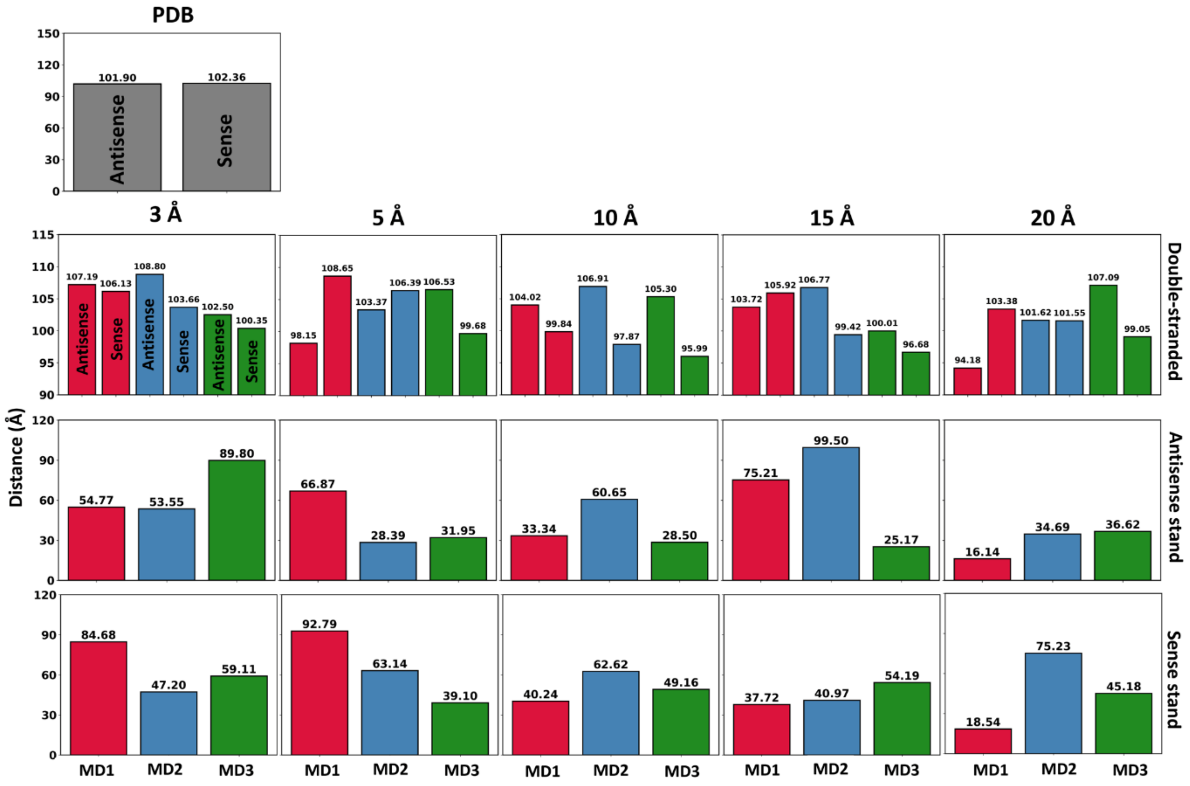


**Figure S12. Terminal residue distance is calculated for three DNA forms (double-stranded, antisense strand, sense strand) across five cell sizes. Grey, red, blue, and green colors represent PDB, RUN1, RUN2, and RUN3.**

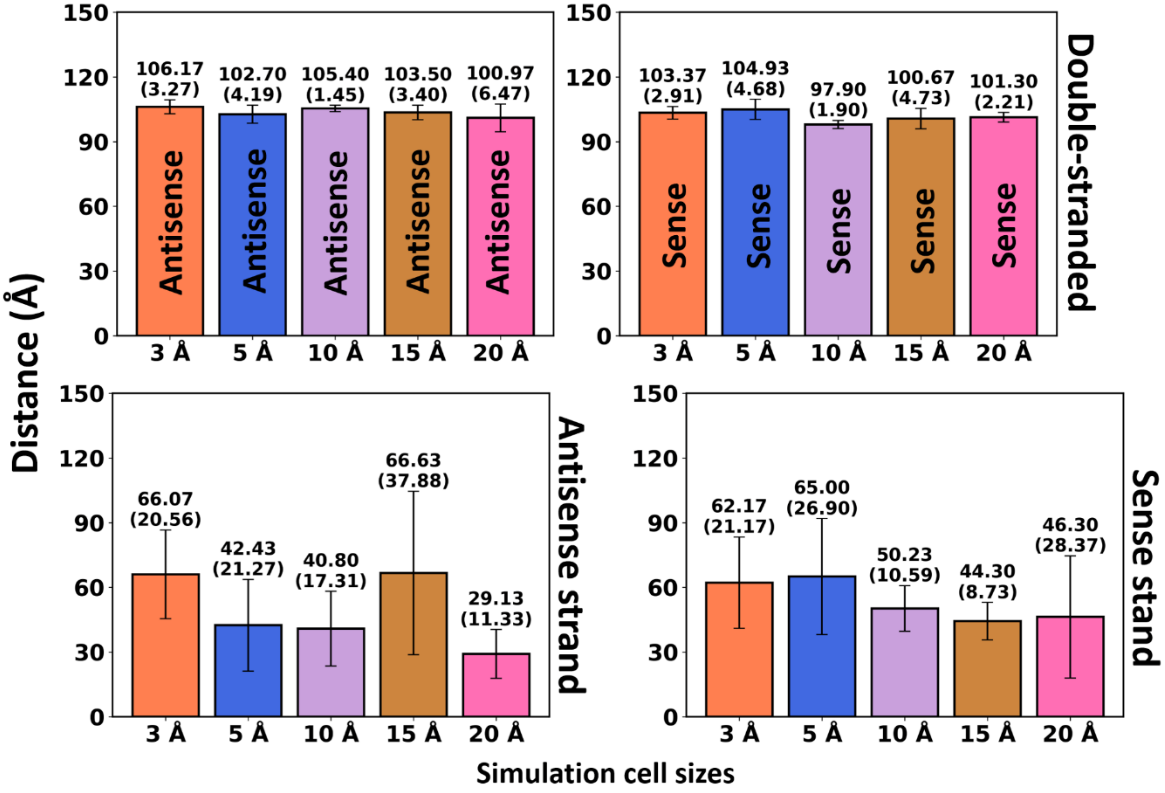


**Figure S13. Average and standard deviation of terminal residue distance for three runs. Calculated for three DNA forms (double-stranded, antisense, sense) across five cell sizes.**

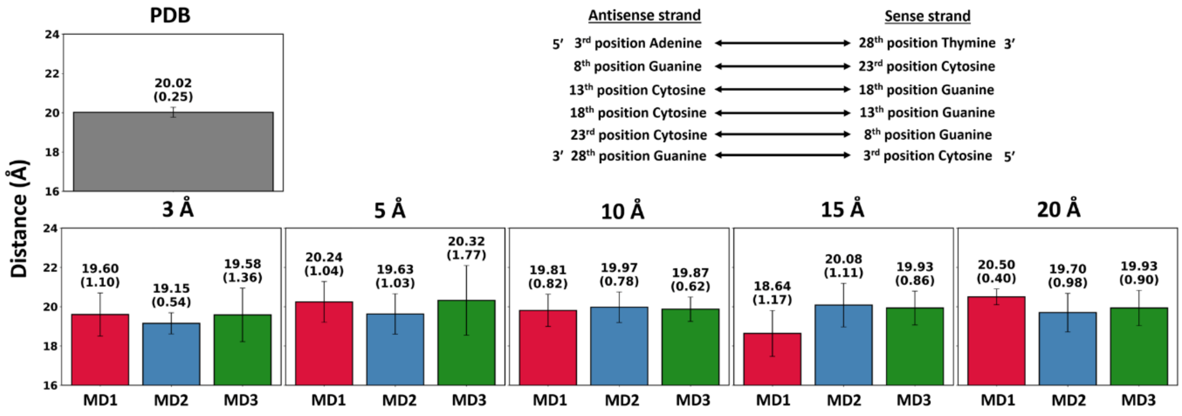


**Figure S14. Distance is calculated for adjacent residues of both strands (antisense and sense) in double-stranded DNA. Residues are selected from 3rd position to 28th position with gap of four residues as mentioned in figure. Terminal residues were not selected because of their direct contact with water lead to greater deviation. Thus, they are not ideal to evaluate diameter parameter (width of DNA) of double-stranded DNA. Grey, red, blue, and green color represent PDB, RUN1, RUN2, and RUN3.**

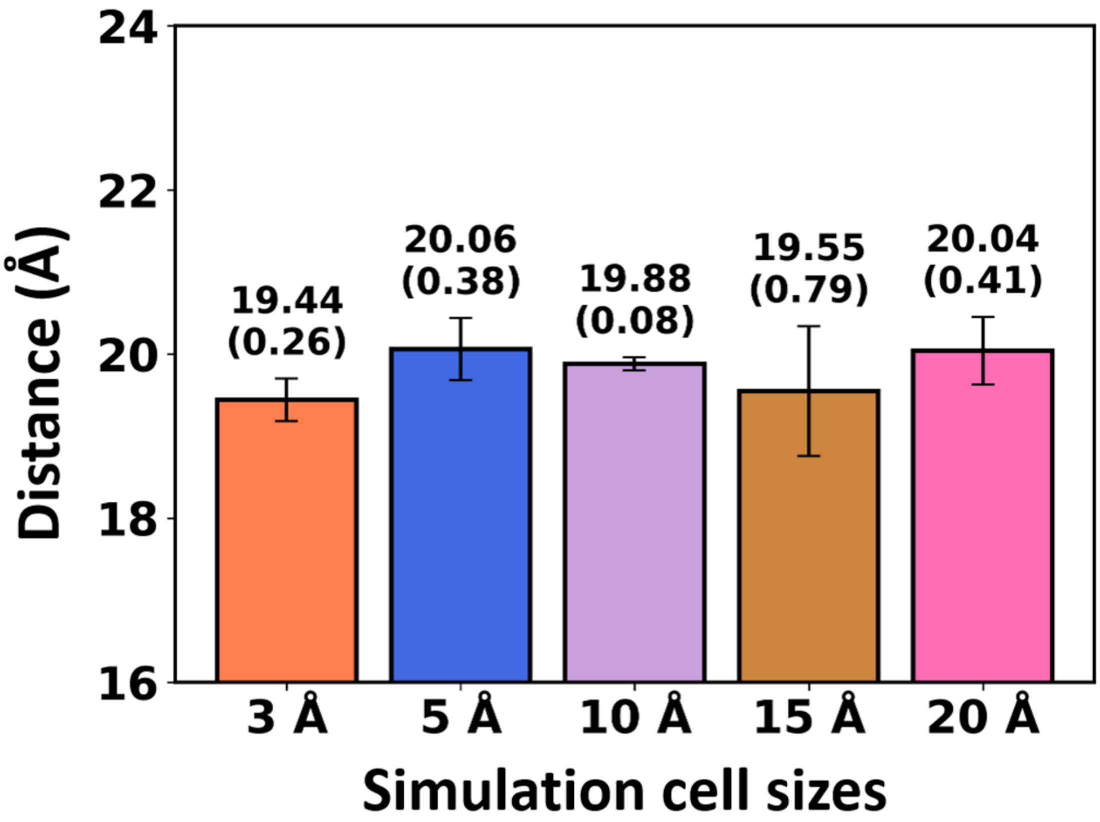


**Figure S15. Average and standard deviation of diameter for three runs. Calculated for double-stranded DNA across five cell sizes. An annotation represents average (top) and stranded deviation (bracket).**
